# DNA replication stress redistributes the histone methyltransferase G9a and activates lineage-specific gene expression

**DOI:** 10.64898/2026.10.06.757090

**Authors:** Gillian E. Oaks, Brian T. Do, Emma M. Jakubicka, Patrick J. Cunniff, Abigail P. Ward, Claire M. Choplick, Samantha A. Nelson, Daniel E. Bauer, Matthew G. Vander Heiden

**Author notes:** Co-first authors (Gillian E. Oaks, Brian T. Do).

## Abstract

Perturbations that cause DNA replication stress can induce lineage-specific changes in gene expression. Specifically, DNA replication stress activates primed loci in euchromatin that are epigenetically silenced, but the mechanism linking stalled replication forks to the activation of distal genes remains unknown. Using reactivation of fetal hemoglobin (HbF) by hydroxyurea in erythroid progenitors as a model, we find that redistribution of the repressive histone methyltransferase G9a from cis-regulatory elements to stalled replication forks increases transcription of the gamma-globin genes *HBG1* and *HBG2*, explaining how hydroxyurea increases HbF expression and benefits patients with sickle cell anemia. In addition to loss of *HBG1/2* silencing, redistribution of G9a also increases expression of a primed, GATA1-driven transcriptional program. G9a and H3K9me2 accumulate at stalled replication forks in a manner dependent on the DNA-binding protein WIZ. Overexpression of G9a partially reduces the impact of G9a redistribution on *HBG1/2* transcript levels. Together, these findings support a model in which stalled replication forks recruit corepressors away from primed genes, thereby facilitating their activation and explaining how DNA replication stress promotes lineage-specific changes in gene expression.

## Introduction

Cells have evolved intricate machinery to maintain the fidelity of gene expression programs and epigenetic state during DNA replication. This requires coordination of multiple processes such as pausing of transcription at replication forks, deposition of new histones, and faithful copying of histone marks^1^. These processes must enable replication of the entire genome and be resilient to stresses that might slow or stall replication forks. However, in pathophysiological settings such as cancer or following treatment with specific drugs, decreased deoxynucleotide availability or other perturbations that impair DNA replication can lead to prolonged DNA replication stress (hereafter referred to as replication stress). Replication stress induces adaptive cellular responses, including those downstream of signaling through the ATR/Chk1 pathway, which can induce expression of genes to enable deoxynucleotide synthesis, DNA fork protection and restart, and DNA repair via pathways that are largely agnostic to cell type^2,3^.

Replication stress can also activate cell type-specific gene expression changes including activation of lineage-dependent transcriptional programs that promote differentiation^4^. Replication stress-induced changes in gene expression state can occur in diverse settings including both transformed and non-transformed hematopoietic cells^4,5^, during pancreatic development^6^, and in mouse embryonic stem cells^7,8^. In these cases, replication stress leads to activation of genomic loci that exist in a primed but epigenetically silenced state characterized by both accessible chromatin and transcriptional repression^9^. The underlying transcription factor landscape of a given cell type appears to dictate the primed genes that are activated when DNA replication is impaired^4^, but the molecular mechanisms which enable the activation of these specific genomic loci in response to replication stress are not well understood.

In sickle cell disease (SCD), increased expression of fetal hemoglobin (HbF) by erythroid progenitor cells in response to hydroxyurea, which inhibits dNTP synthesis and induces replication stress^10^, is a clinically relevant example of gene reactivation following replication stress. However, despite decades of hydroxyurea being used as a first-line treatment for SCD, the mechanism by which it induces HbF expression remains unclear^10,11^. Functional hemoglobin is a tetrameric molecule containing two α-globin subunits and two subunits of either γ-globin (yielding fetal hemoglobin, or HbF) or β-globin (yielding adult hemoglobin, or HbA), which are encoded by adjacent genes which reside within the β-globin locus on chromosome 11: γ-globin by the *HBG1* and *HBG2* genes (hereafter referred to as just *HBG*) and β-globin by the *HBB* gene^12^. Humans predominantly express HbF until approximately six months post-conception, after which erythroid precursors gradually silence *HBG* while upregulating *HBB*, leading to a switch from HbF to HbA expression^13^. This switch is enforced by sequence-specific repressors such as BCL11A^14^ and LRF/ZBTB7A^15^, as well as corepressors such as NuRD, LSD1/CoREST, and the G9a/GLP complex^16^, that bind to regulatory elements surrounding *HBG*. Reactivating *HBG* to increase HbF levels is the primary therapeutic strategy in SCD, in which β-globin is mutated, and in beta-thalassemia, in which β-globin production is compromised. While treatment with hydroxyurea can confer partial reactivation of HbF^10^, stronger reactivation has been achieved through recently FDA-approved gene therapy to disrupt BCL11A expression in hematopoietic stem and progenitor cells^17^ and molecular glue compounds currently in clinical trials that degrade LRF/ZBTB7A^18^ or WIZ^19^, which recruits the G9a/GLP complex to *HBG*^20^. Nevertheless, how hydroxyurea reactivates HbF and whether it modulates the activity of epigenetic machinery at the *HBG* locus is not known.

Here, we find that replication stress is accompanied by redistribution of corepressors from the *HBG* locus to stalled replication forks. We observe loss of the repressive histone marks H3K9me2 and H3K9me3, as well as reduced binding of the G9a/WIZ corepressor complex that deposits H3K9me2, at the globin locus without changes in chromatin accessibility. This leads to transcriptional activation of both *HBG* and *HBB*, with a relative increase in HbF expression. Loss of G9a/WIZ binding and H3K9me2 occupancy also occurs genome-wide in response to replication stress. In erythroid progenitor cells, this leads to derepression of primed regulatory loci, ultimately inducing a GATA1/2-dominated transcriptional program. Proximity ligation and immunoprecipitation of proteins bound to newly replicated DNA demonstrate redistribution of G9a to sites of replication stress, and increased expression of G9a partially suppresses *HBG* activation by replication stress. These data argue that stalled replication forks sequester the G9a-containing corepressor complex from target loci across the genome, explaining how hydroxyurea induces *HBG* expression to increase HbF levels in erythroid cells. More broadly, these data provide evidence that stalled replication forks sequester the G9a-containing corepressor complex from target loci across the genome and suggest a mechanism for how replication stress activates primed transcriptional programs to induce lineage-specific changes in cell state.

## Results

### DNA replication stress activates gene expression at the globin locus, increasing relative HbF expression

HUDEP-2 immortalized erythroid progenitor cells are a model for studying and modulating expression of genes at the globin locus^21^. Treatment with hydroxyurea for 48 hours led to a dose-dependent increase in the percentage of HbF-expressing cells as measured by intracellular flow cytometry **(Fig. 1A)**. Hydroxyurea treatment also led to dose-dependent increases in levels of CHEK1 phosphorylation at Ser345 (p-Chk1), a marker of replication stress **(Fig. 1B)**. Of note, hydroxyurea treatment increased transcription of both *HBG* and *HBB* as measured by RNA sequencing of nascent transcripts **(Fig. 1C)**. To test whether hydroxyurea induces HbF expression by causing replication stress, we tested a variety of pharmacologic agents that cause replication stress via orthogonal mechanisms, including aphidicolin, which directly inhibits DNA polymerases^22^, camptothecin and etoposide, which inhibit topoisomerases^23^, decitabine, which traps DNA methyltransferases on chromatin^24^, and methotrexate, which inhibits purine synthesis^25^. Each perturbation induced HbF expression, including at doses that also increase p-Chk1 without substantially impairing viability or increasing phosphorylation of Thr68 on CHEK2 (p-Chk2), a marker of DNA double-strand breaks (**Figs. 1D-E, S1A-B**). Taken together, these results suggest that diverse perturbations that cause replication stress can increase HbF expression in HUDEP-2 cells, independent of causing widespread DNA damage.

**Figure 1:**
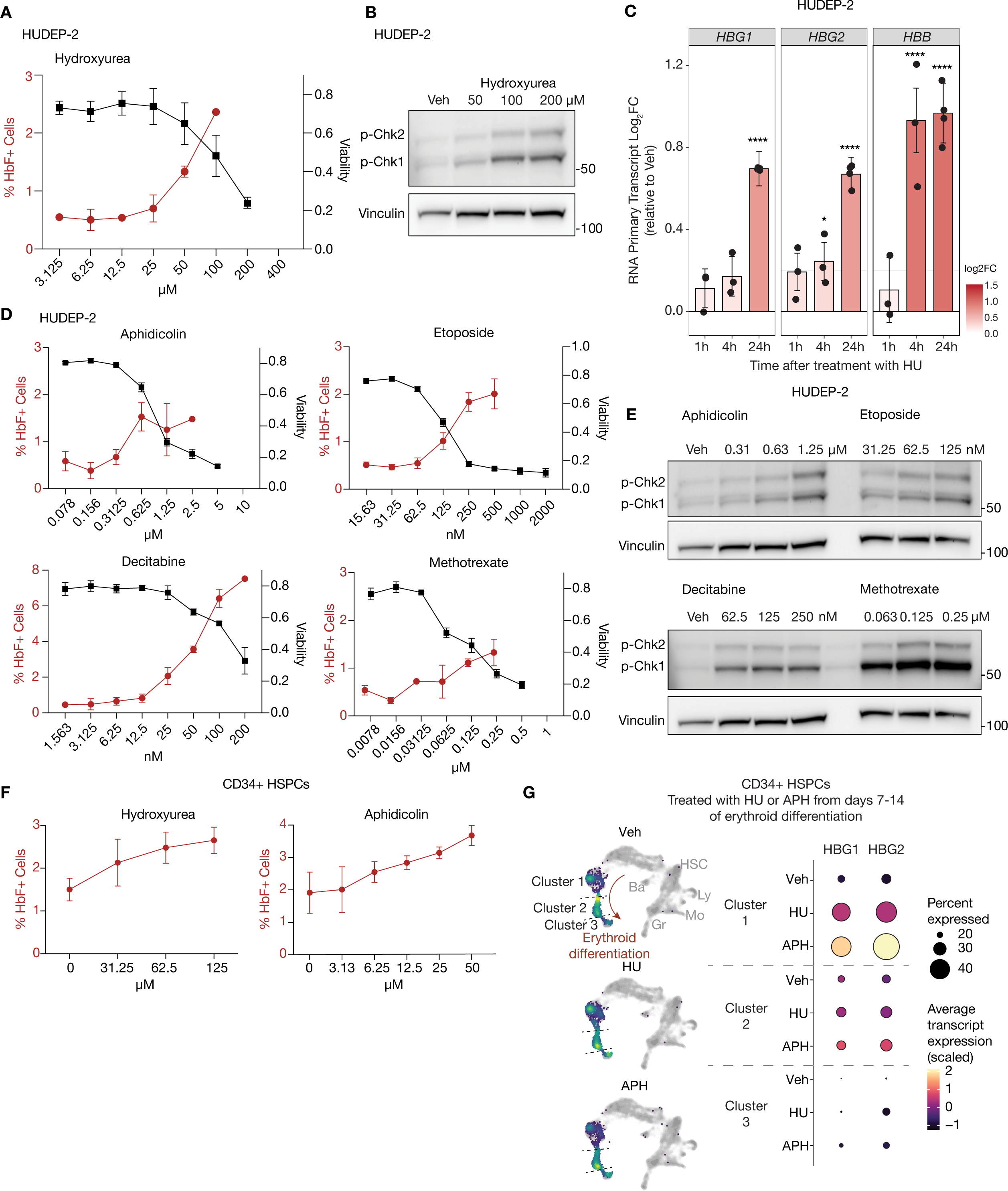
DNA replication stress induces gamma-globin expression in erythroid progenitors. **(A)** Dose-response curve of fetal hemoglobin induction (HbF, red, left axis) and cell viability (black, right axis) in HUDEP-2 cells treated with increasing concentrations of hydroxyurea (HU) (n=3). **(B)** Representative immunoblots of HUDEP-2 cells treated for 48 hr with the indicated doses of HU. Phosphorylation of CHEK1 at Ser345 (p-Chk1) is shown as a marker of DNA replication stress and phosphorylation of CHEK2 at Thr68 (p-Chk2) as a marker of DNA damage. Vinculin serves as a loading control. **(C)** Nascent RNA expression analysis of globin genes (*HBG1*, *HBG2*, and *HBB*) in HUDEP-2 cells at the indicated times following treatment with HU. Bar plots show the mean log2 fold change (log_2_FC) of primary transcript levels relative to vehicle at the indicated time points after treatment. Asterisks denote statistically significant (p < 0.00001) changes compared to vehicle (n=4). **(D)** Dose-response curve of HbF induction (red, left axis) and cell viability (black, right axis) in HUDEP-2 cells treated with increasing concentrations of aphidicolin, etoposide, decitabine, and methotrexate as indicated (n=3). **(E)** Immunoblot analysis of HUDEP-2 cells treated as described in (D) for p-Chk1, p-Chk2, and Vinculin. **(F)** Dose-response curves of HbF induction in primary human CD34^+^ hematopoietic stem and progenitor cells (HSPCs) treated with increasing concentrations of hydroxyurea or aphidicolin (n=3). **(G)** Single-cell RNA-seq analysis of CD34+ HSPCs treated with vehicle (Veh), HU, or aphidicolin (APH) from days 7–14 of erythroid differentiation and analyzed on day 14. (Left) UMAP projections showing density plots of cells from each treatment condition, overlaid on the reference hematopoietic differentiation landscape (gray). Clusters 1-3 are labeled. (Right) Dot plots showing *HBG1* and *HBG2* expression within erythroid clusters 1-3 for each treatment condition. Dot size represents the percentage of cells expressing each transcript and color indicates scaled, log-normalized average expression.

To determine whether this phenotype of replication stress inducing HbF expression extends to non-immortalized hematopoietic cells, we examined HbF expression in primary human CD34^+^ hematopoietic stem and progenitor cells (HSPCs) under conditions that cause erythroid differentiation. Treatment with hydroxyurea, aphidicolin, or decitabine for 48 hours at day 7 of erythroid expansion led to increased HbF and p-Chk1 as measured by flow cytometry **(Figs. 1F, S1C-E**), suggesting that replication stress can also promote HbF expression in HSPCs. Replication stress also induces differentiation markers in several hematopoietic cell models^26^. Consistent with this, we observed increased expression of the erythroid differentiation markers CD71 and CD235a in HUDEP-2 cells following hydroxyurea exposure **(Fig. S1F)**. Reanalysis of a previously published single-cell RNA sequencing dataset of CD34^+^ HSPCs treated with hydroxyurea and aphidicolin for seven days starting at day 7 of erythroid differentiation demonstrated that increased *HBG* expression was not attributable to a change in differentiation state **(Figs. 1G, S1G)**. Thus, induction of HbF by hydroxyurea and other inducers of replication stress cannot solely be attributed to differentiation.

### A live-cell reporter of HbF enables study of globin activation dynamics

To enable investigation of *HBG* expression dynamics in live cells, we generated HUDEP-2 cells with an mCherry fluorescent reporter co-expressed from the endogenous *HBG2* locus^27^ **(Fig. S2A)**. Consistent with results assessing HbF expression in response to replication stress, HBG2-mCherry cells showed increased mCherry expression over time in response to hydroxyurea or decitabine treatment **(Fig. S2B)**, and mCherry expression correlated with HbF protein expression **(Fig. S2C)**. To further verify this model, we sorted populations of cells with low, medium, and high mCherry fluorescence and found that the amount of mCherry fluorescence correlates with *HBG* RNA transcript levels, but not with levels of *HBB* RNA transcripts **(Fig. S2D)**. As previously described^28^, individual HUDEP-2 cells vary widely in HbF expression. Thus, to reduce variation in the model system, we sorted individual clones with varying levels of baseline mCherry fluorescence **(Fig. S2E)**. Higher baseline expression of HbF was associated with HbF induction at lower doses of hydroxyurea and aphidicolin, despite hydroxyurea and aphidicolin having similar effects on cell viability in all clones **(Fig. S2F)**. These data further confirm that HbF reactivation in response to replication stress is independent of changes in viability and suggest that the degree of reactivation may be influenced by preserved, cell-intrinsic features.

It is unclear whether cells with HU-induced increases in HbF expression arise from a subpopulation of erythroid progenitors that express some HbF at baseline, or whether HU can induce expression of HbF in cells that predominantly express HbA. To address this, we analyzed the behavior of mCherry-low cells (HbF low and HbA predominant, referred to as A cells) and mCherry-high cells (higher baseline HbF expression, referred to as F cells) sorted from a bulk population (**Fig. S2G**). Post-sort examination revealed that A cells can spontaneously give rise to F cell progeny at low rates and that both hydroxyurea and decitabine treatment (as well as pretreatment with these two compounds) can increase the frequency of F cell generation (**Fig. S2H, left**). We also observed that the frequency of reversion from F cells to A cells was decreased when hydroxyurea treatment was continued after the sort **(Fig. S2H, right)**. Interestingly, past treatment with decitabine, but not hydroxyurea, decreased the frequency of reversion from the F state to the A state, possibly because of the additional DNA hypomethylating properties of decitabine^24,29^. To test whether these observations might be explained by a heritable cell state, we single-cell cloned A cells, grew them into small colonies of 100-1000 cells, and then split these colonies into three sister colonies, one of which was treated with vehicle and two of which were treated with hydroxyurea for 3 days **(Fig. S2I)**. We found that the hydroxyurea-treated sister colonies had highly correlated degrees of mCherry induction that were not significantly associated with the mCherry percentage in their vehicle-treated sister colony **(Fig. S2J-K)**. These data indicate that heritable factors that persist over at least 7-10 cell doublings primarily influence the degree to which A cells can reactivate HbF in response to hydroxyurea treatment.

### Induction of HbF is independent of HRI and ATR signaling

We next examined whether known stress response mechanisms might connect replication stress to *HBG* reactivation. Knockdown of HRI, a kinase activated by mitochondrial stress to induce the integrated stress response, has previously been shown to activate HbF^30,31^. Using shRNA knockdown of HRI in HUDEP-2 cells, we confirmed that knockdown of HRI increased baseline HbF levels but also found that treatment with replication stress inducers further increased HbF, implying that replication stress can increase HbF independently of HRI (**Fig. S3A-B)**. The ATR kinase mediates many cellular responses to replication stress^3^. To assess a requirement for ATR in replication stress-induced HbF expression, we treated HUDEP-2 mCherry cells with AZ20, an inhibitor of ATR and focused on early timepoints to avoid cell death and minimize secondary effects of ATR inhibition, which itself can cause replication stress^32^. We found that ATR inhibition did not prevent either hydroxyurea- or aphidicolin-driven increases in mCherry signal. Additionally, ATR inhibition with 1µM AZ20 on its own led to a small increase in mCherry signal, consistent with the replication stress observed at this dose **(Fig. S3C-E)**. Thus, transcriptional induction of *HBG* expression does not require ATR signaling.

### Hydroxyurea treatment decreases repressive chromatin marks and corepressor binding genome-wide, including at the globin locus

To explore the epigenetic consequences of hydroxyurea treatment on the globin locus, as well as genome-wide, we used cleavage under targeting and release using nuclease (CUT&RUN)^33^ to map the repressive histone modifications H3K9me2, H3K9me3, and H3K27me3 and the activating modifications H3K9ac and H3K27ac in HUDEP-2 *HBG-*mCherry cells that exhibit about 5% mCherry+ cells at baseline (Clone A, **Fig. S2E)**. At baseline, the globin locus contained histone modifications associated with both active gene expression and transcriptional repression **(Fig. 2A)**. Specifically, we observed high levels of activating marks, including H3K9ac and H3K27ac at the *HBG1*, *HBG2,* and *HBB* promoters, in intervening regions such as the *HBBP1* pseudogene, and in the locus control region (LCR) while the repressive mark H3K27me3 was almost entirely absent **(Fig. 2A)**. However, the repressive marks H3K9me2 and H3K9me3 were observed at many places within the globin locus: H3K9me2 was present at the LCR and *HBG* and *HBB* gene bodies, and H3K9me3 was present between *HBG* and *HBE* **(Fig. 2A)**. Upon hydroxyurea treatment, we observed a concordant decrease in H3K9me2 signal and increase in H3K9ac signal at both the *HBB* and *HBG* loci and the LCR. We also observed reduced H3K9me3 flanking the globin locus. H3K27ac signal was markedly decreased at the *HBB* locus, but relatively unchanged at the *HBG* loci **(Fig. 2A)**, in accordance with a shift to increased *HBG* expression.

**Figure 2:**
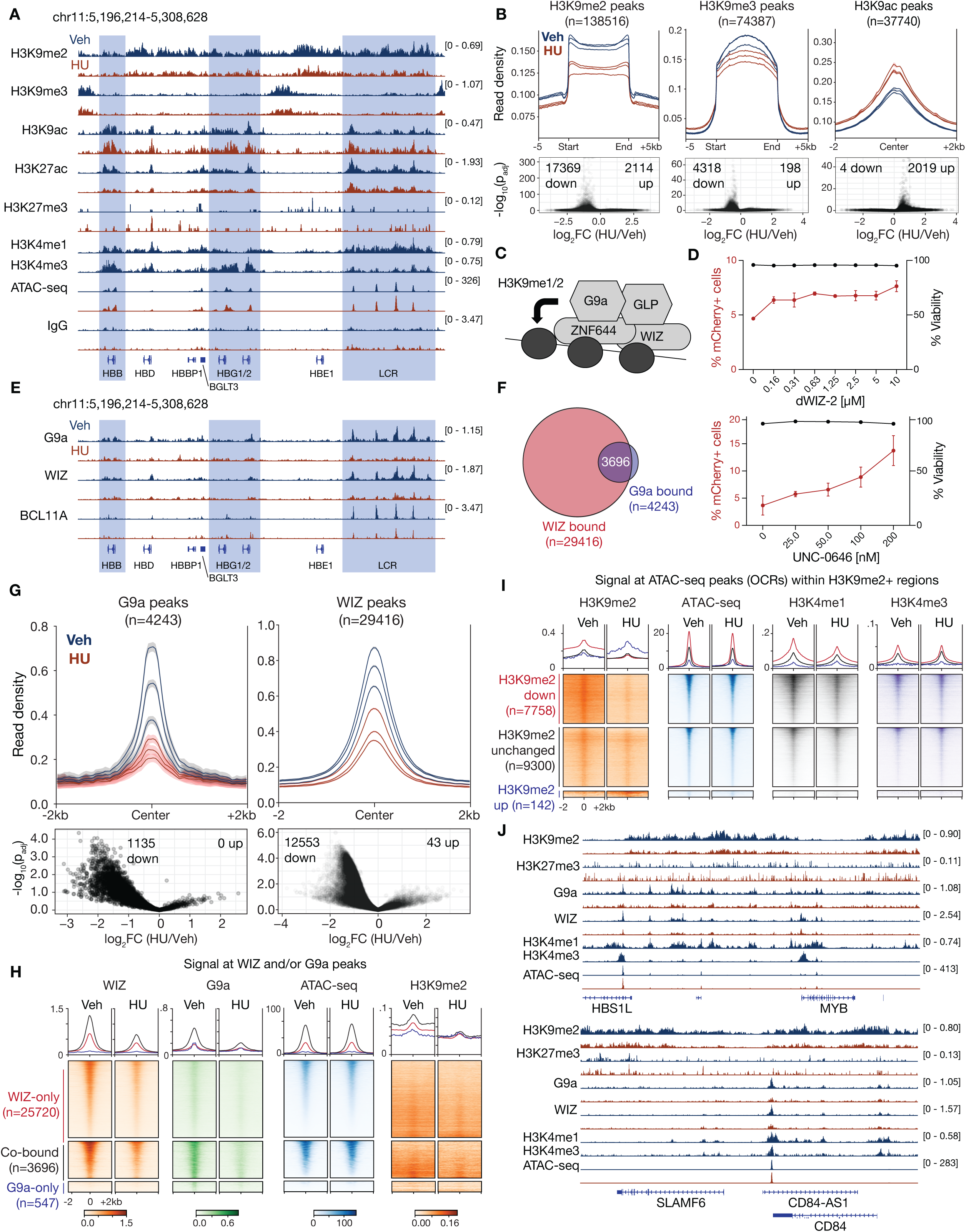
Hydroxyurea decreases H3K9me2 and binding of G9a, WIZ and BCL11A at the globin locus and throughout the genome. **(A)** Genome browser tracks showing CUT&RUN profiles for H3K9me2, H3K9me3, H3K9ac, H3K27ac, H3K27me3, H3K4me1, H3K4me3, and IgG control as well as ATAC-seq signal at the globin locus under vehicle-treated (Veh, blue) and hydroxyurea-treated (HU, red) conditions. Highlighted regions denote globin gene clusters and the globin locus control region (LCR) as indicated. **(B)** Metagene profiles (top) and corresponding differential analyses (bottom) for H3K9me2, H3K9me3, and H3K9ac peaks in Veh and HU conditions. Read density is plotted across peak centers ±5 kb (H3K9me2/3) or ±2 kb (H3K9ac) and corresponding volcano plots show log_2_ fold change of HU relative to Veh (log_2_FC, HU/Veh) versus –log_10_ adjusted p-value (p_adj_), with numbers of significantly decreased (down) and increased (up) peaks indicated. **(C)** Schematic of the G9a-GLP-WIZ-ZNF644 complex associating with DNA and depositing H3K9me1/2 histone marks. **(D)** Dose-response analysis of mCherry expression (red, left axis) and cell viability (black, right axis) in HUDEP-2 HBG-mCherry cells treated with increasing concentrations of dWIZ-2 (top) and UNC-0646 (bottom) (n=3). **(E)** Genome browser tracks showing CUT&RUN profiles for G9a, WIZ, and BCL11A at the globin locus under Veh (blue) and HU (red) conditions. Highlighted regions denote globin gene clusters and the globin locus control region (LCR) as indicated. **(F)** Venn diagram showing overlap between WIZ-bound and G9a-bound peaks as indicated. The number of peaks for each protein is denoted. **(G)** Metaplots (top) and differential analyses (bottom) of G9a and WIZ CUT&RUN peaks under Veh and HU-treated conditions. Read density is plotted across peak centers ±2 kb and corresponding volcano plots show log_2_ fold change of HU relative to Veh (log_2_FC (HU/Veh)) versus –log_10_ of adjusted p-value (-log_10_(p_adj_)), with numbers of significantly decreased (down) and increased (up) peaks indicated. **(H)** Metaplots (top) and heatmaps (bottom) showing signal intensity for WIZ CUT&RUN, G9a CUT&RUN, ATAC-seq, and H3K9me2 CUT&RUN at three classes of regions: WIZ-only peaks, WIZ/G9a co-bound peaks, and G9a-only peaks as indicated. Signals are plotted ±2 kb from peak centers under Veh and HU-treated conditions. **(I)** Metaplots (top) and heatmaps (bottom) for H3K9me2 CUT&RUN, ATAC-seq, H3K4me1 CUT&RUN, and H3K4me3 CUT&RUN at OCRs within H3K9me2+ regions as indicated. Regions are grouped based on changes in H3K9me2 levels upon HU treatment (down, unchanged, up), under both Veh and HU-treated conditions. **(J)** CUT&RUN profiles showing H3K9me2, H3K27me3, G9a, WIZ, H3K4me1, and H3K4me3 as well as ATAC-seq signal during vehicle treatment (blue) or HU treatment (red) for representative genes that show loss of repressive histone marks and euchromatic corepressors in response to HU treatment.

In addition to the effects of hydroxyurea on the globin locus, we observed reduced H3K9me2 and H3K9me3 signal with slight increases in H3K9ac across the genome **(Fig. 2B)**. These same patterns were observed upon hydroxyurea treatment in HUDEP-2 mCherry cells with a higher baseline level of mCherry-positive cells (20-30%) (Clone C, **Figs. S2E**, **S4A)**. For both clones, we also confirmed that the changes observed upon hydroxyurea treatment were not due to differences in pulldown efficiency by normalizing CUT&RUN signal against spike-ins of a known quantity of mouse nuclei **(Figs. S4B-C, S4E).** Taken together, these data suggest that reactivation of *HBG* by hydroxyurea is associated with reduced H3K9me2 deposition at the globin locus, including at *HBG*.

While H3K9me2 is often observed as an intermediate leading to H3K9me3 during formation of heterochromatin, H3K9me2 can also be found at loci that are actively silenced within euchromatin^34^. At these loci, H3K9me2 is primarily deposited by the G9a/GLP methyltransferase complex, a four-subunit complex consisting of the methyltransferases G9a and GLP together with the DNA-binding proteins WIZ and ZNF644^34,35^ **(Fig. 2C)**. As a corepressor, this complex is recruited to genes by different repressive TFs in different cellular contexts^36^. Chemical inhibition of G9a methyltransferase activity using the compound UNC-0646 or degradation of WIZ using the compound dWIZ-2 have been previously shown to reduce H3K9me2 deposition at the globin locus^16,19^. We therefore hypothesized that hydroxyurea suppresses H3K9me2 signal at the globin locus by modulating activity of the G9a/GLP complex, and that disruption of the complex would phenocopy the effects of hydroxyurea exposure on HbF expression. Indeed, both UNC-0646 and dWIZ-2 exposure increased mCherry expression in HBG2-mCherry HUDEP-2 cells **(Figs. 2D, S4F)**. Correspondingly, CUT&RUN of G9a and WIZ demonstrated reduced binding of these factors across the globin locus upon hydroxyurea treatment, with the largest changes observed at the LCR and at the *HBG1* and *HBG2* promoters **(Figs. 2E, S4D)**. Moreover, hydroxyurea treatment decreased CUT&RUN signal for BCL11A **(Fig. 2E)** at *HBG* and within the LCR^16^. These observations suggest that hydroxyurea exposure reduces binding of key repressive machinery across the globin locus and specifically at *HBG*.

We next assessed genome-wide localization of WIZ and G9a in response to hydroxyurea treatment. In untreated cells, WIZ was found at ∼29,000 peaks and G9a was found at ∼4,200 peaks, including ∼3,800 overlapping peaks **(Fig. 2F)**. While changes in WIZ and G9a binding after 24 hours of hydroxyurea exposure were correlated, hydroxyurea treatment led to a larger decrease in G9a binding (∼80% reduction) than WIZ binding (∼60% reduction), possibly because WIZ directly binds DNA^20^ **(Figs. 2G, S4G)**. At cis-regulatory elements co-occupied by WIZ and G9a, H3K9me2 levels were reduced, and H3K9ac levels were increased following hydroxyurea exposure, with no significant change in H3K9me3 **(Figs. 2H, S4H)**. The degree of decrease in H3K9me2 signal upon hydroxyurea treatment was also associated with proximity to WIZ occupancy **(Fig. S4I)**. BCL11A also colocalized with WIZ and G9a and was evicted from WIZ/G9a bound loci following exposure to hydroxyurea **(Figs. S4J-K)**. Thus, hydroxyurea exposure is associated with a genome-wide decrease in H3K9me2 as well as reduced G9a, WIZ, and BCL11A binding.

### Loss of repressive marks at primed loci occurs without increases in chromatin accessibility

The selective loss of the primed chromatin mark H3K9me2 and decreased binding of the euchromatic corepressors G9a and WIZ genome-wide suggested that hydroxyurea may specifically act at primed sites: euchromatic cis-regulatory elements that are transcriptionally repressed. To test this, we first measured chromatin accessibility using Assay for Transposase-Accessible Chromatin with sequencing (ATAC-seq)^37^ in HBG2-mCherry HUDEP-2 cells with and without hydroxyurea treatment **(Figs. 2A, S5A)**. These experiments were also performed in sorted populations of A and F cells to assess whether these populations differed in chromatin accessibility, especially within the globin locus. We found that the *HBG* promoters were accessible in both A and F cells, as well as before and after exposure to hydroxyurea treatment. Genome-wide chromatin accessibility was minimally altered between A and F cells and following hydroxyurea treatment. Only ∼1% of open chromatin regions (OCRs) showed statistically significant changes in accessibility, and differences between hydroxyurea and vehicle-treated accessibility programs were largely driven by erythroid differentiation-related TF motifs whereas differences between A and F cells were related to NF-Y and AP-1 motifs **(Figs. S5B-C)**. We next categorized open chromatin regions (OCRs) based on whether they gained or lost H3K9me2 upon hydroxyurea treatment. We found that regions of chromatin that lost H3K9me2 signal upon hydroxyurea treatment were characterized by greater chromatin accessibility at baseline compared to regions that had unchanged or increased H3K9me2 signal, supporting the hypothesis that hydroxyurea causes loss of H3K9me2 at primed loci **(Fig. 2I)**. These loci also did not gain further accessibility upon hydroxyurea treatment, consistent with previous observations that cell state changes upon replication stress are not associated with large changes in chromatin accessibility^4^.

To further characterize the primed loci that lose H3K9me2, we performed CUT&RUN for the enhancer-associated mark H3K4me1, the promoter-associated mark H3K4me3, and the active-chromatin mark H3K27ac. To our surprise, we identified all three marks at *HBG1/2* promoters at baseline, consistent with a primed but actively repressed state **(Fig. 2A)**. Furthermore, these marks were also enriched at OCRs that lost H3K9me2 (“H3K9me2-down”) compared to OCRs that had unchanged or increased H3K9me2 following hydroxyurea treatment **(Figs. 2I, S5D)**. Of the other marks analyzed, only the activating mark H3K9ac increased at H3K9me2-down OCRs following hydroxyurea treatment, suggesting that loss of corepressor occupancy is a trigger for locus derepression **(Fig. S5D)**. Additionally, upon hydroxyurea treatment, we observed that loss of H3K9me2, G9a, and WIZ binding occurs near hematopoietic lineage-related genes such as *MYB*, *KIT*, *RAG1/2*, and *SLAMF6* **(Figs. 2J, S5E)**.

Another repressive mark associated with active silencing is H3K27me3, which is deposited by polycomb repressive complex 2 (PRC2). We did not detect H3K27me3 at the *HBG* loci **(Figs. 2A, S4A)**. Genome-wide, hydroxyurea treatment led to some loss of H3K27me3, although only 439 regions significantly lost H3K27me3 compared to 17369 regions that significantly lost H3K9me2 **(Fig. S5F)**. OCRs that lost H3K27me3 (“H3K27me3-down”) in response to hydroxyurea treatment had fewer signs of active transcription (lower H3K4me1, H3K27ac, and ATAC-seq signal) relative to OCRs that lost H3K9me2 (“H3K9me2-down”) **(Fig. 2I, S5G)**, emphasizing that H3K9me2-down regions exist in a less repressed, more primed state than that associated with classic heterochromatin. Interestingly, WIZ binding at baseline was similar in both sets of OCRs, although G9a and BCL11A were more highly bound at H3K9me2-down OCRs compared to H3K27me3-down OCRs **(Fig. S5H)**. Notably, regions that lost H3K27me3 tended to be separate from those that lost H3K9me2 and were enriched for gene ontology categories and transcription factor motifs associated with non-hematopoietic lineages such as craniofacial development **(Fig. S5I-L)**. Taken together, these data suggest hydroxyurea treatment is also associated with reactivation of a small set of non-erythroid loci marked by H3K27me3.

### In erythroid progenitor cells, loss of corepressor binding is associated with derepression of primed erythroid lineage genes

Consistent with an established role for G9a as a euchromatic corepressor^35^, approximately 95% of genes located near regions co-bound by G9a and WIZ were associated with ATAC-seq positive regions **(Fig. S6A)**, suggesting that loss of G9a and H3K9me2 at these loci may have transcriptional consequences. We first performed transcription factor motif analysis using HOMER^37^ on G9a-bound regions, WIZ-bound regions, and regions that significantly lost H3K9me2 following hydroxyurea treatment. Across all three analyses, GATA family transcription factor motifs were among the most highly enriched **(Figs. 3A-C)**. As there is substantial overlap between G9a and BCL11A CUT&RUN peaks **(Fig. S4J)**, we also analyzed the top motifs at BCL11A peaks and again found GATA to be the most enriched motif **(Fig. S6B)**. Furthermore, Enrichr analysis^38–40^ of genes associated with H3K9me2-down OCRs revealed strong enrichment for GATA1 and GATA2 regulation, as well as pathways involved in hematopoietic development **(Figs. 3D-E)**. These findings suggest that, in HUDEP-2 erythroid progenitor cells, G9a and WIZ preferentially repress a GATA1/2 transcriptional program.

**Figure 3:**
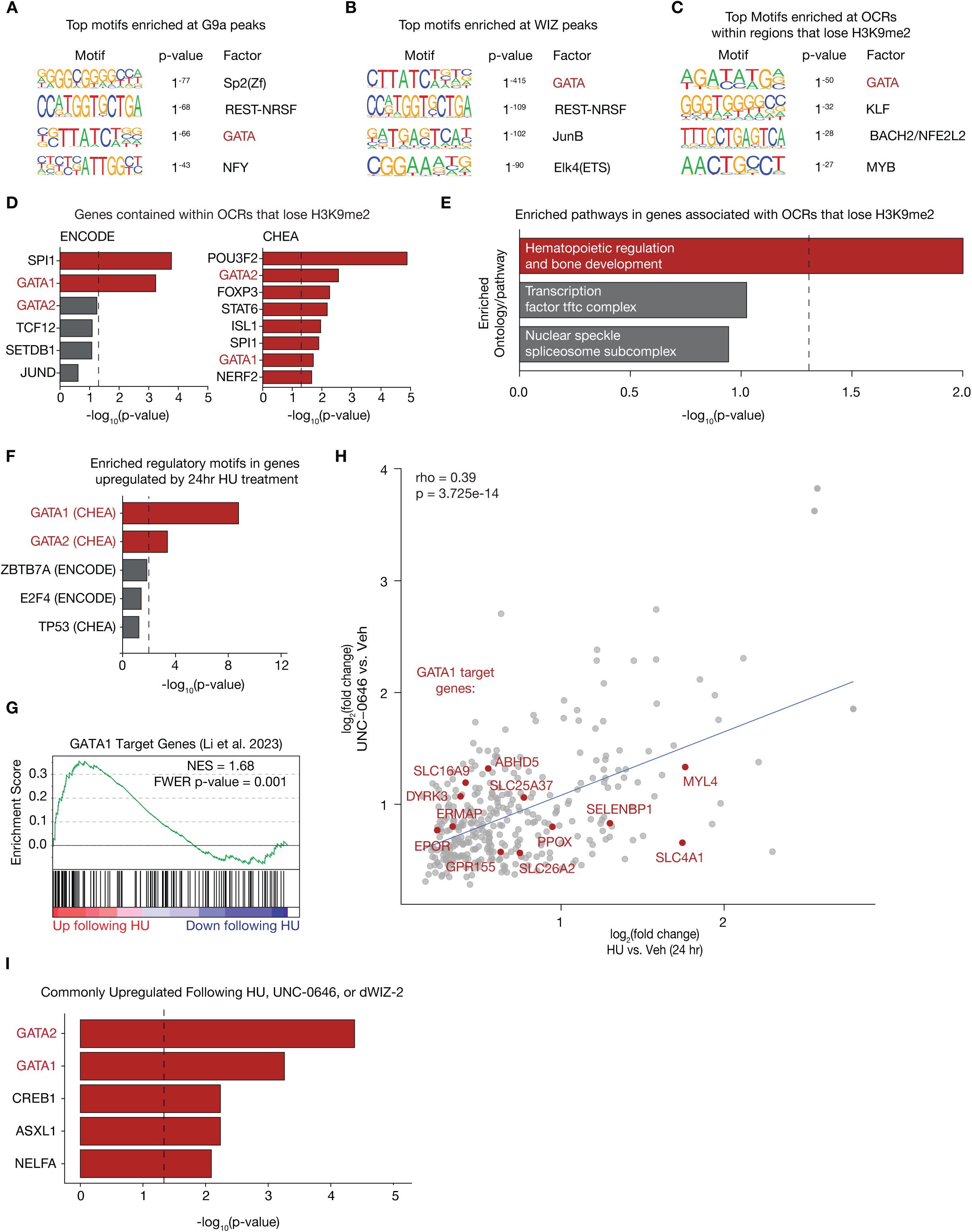
Hydroxyurea treatment leads to upregulation of GATA1/2-regulated genes. **(A)** Transcription factor motif enrichment analysis at G9a-bound peaks. P-values indicate significance of enrichment compared to a background set of peaks as calculated by HOMER. **(B)** Transcription factor motif enrichment analysis at WIZ-bound peaks. P-values indicate significance of enrichment compared to a background set of peaks as calculated by HOMER. **(C)** Transcription factor motif enrichment analysis at ATAC-seq peaks (OCRs) within regions that lose H3K9me2 upon treatment with HU. P-values indicate significance of enrichment compared to a background set of peaks as calculated by HOMER. **(D)** Transcription factor enrichment analysis (CHEA and ENCODE) for TFs associated with genes that lie within OCRs that lose H3K9me2 upon HU treatment. Bar plots show top enriched transcription factors, ranked by –log_10_ of adjusted p-value. Dashed line represents p=0.05. **(E)** Pathway enrichment analysis for genes whose transcription start sites lie within OCRs that lose H3K9me2 upon HU treatment. **(F)** Transcription factor enrichment analysis for TFs associated with genes upregulated following 24 hr HU treatment. Bar plot shows the top enriched transcription factor binding signatures from ENCODE and CHEA datasets, ranked by –log_10_ of adjusted p-value. **(G)** Gene set enrichment analysis (GSEA) for GATA1 target genes identified in a previously published dataset^41^ upon 24 hr HU treatment. NES and FWER p-value are indicated. **(H)** Scatterplot of genes upregulated by HU treatment, with log_2_ fold change in HU relative to vehicle shown on the x-axis and log_2_ fold change in UNC-0646 relative to vehicle on the y-axis. An upregulated subset of the GATA1 target genes used in (G) are labeled and Spearman’s correlation coefficient and associated p-value are indicated. **(I)** Transcription factor enrichment analysis of genes upregulated following HU, UNC-0646, and dWIZ-2 treatment. Bar plot shows the top enriched transcription factor binding signatures from ENCODE and CHEA datasets, ranked by –log_10_ of adjusted p-value.

We next asked whether a GATA1/2-regulated transcriptional program is activated by replication stress. We performed nuclear RNA-seq after 24 hours of hydroxyurea treatment and found that genes bearing GATA1/2 regulatory motifs were the most differentially upregulated gene set **(Fig. 3F)**. To orthogonally validate this observation, we derived a GATA1 target gene set using publicly available RNA-seq data from GATA1 CRISPR-Cas9 knockdown in HUDEP-1 erythroid progenitor cells^41^. Gene set enrichment analysis (GSEA)^42,43^ confirmed significant upregulation of GATA1 target genes following hydroxyurea exposure **(Fig. 3G)**. In previously published single-cell RNA sequencing data from human CD34^+^ HSPCs, hydroxyurea treatment led to accelerated erythroid differentiation^4^, consistent with GATA1 being the dominant driver of erythroid differentiation^44^.

Next, to evaluate the degree to which hydroxyurea exposure phenocopies the transcriptional effect of direct G9a inhibition, we performed nuclear RNA-seq in HUDEP-2 cells following treatment with dWIZ-2^19^ (which degrades WIZ) or UNC-0646^45^ (a catalytic inhibitor of G9a) **(Fig. S6C)**. Among genes upregulated following both UNC-0646 and dWIZ-2 treatment, GATA1 and GATA2 were the most highly enriched regulatory motifs **(Fig. S6D)**. Moreover, the most differentially upregulated genes following hydroxyurea treatment and both UNC-0646 and dWIZ-2 were significantly correlated in their expression change and included known erythroid and GATA1 target genes such as *EPOR*, *ERMAP*, *SLC4A1*, and *SLC25A37* **(Fig. 3H, S6E)**. These genes were also enriched for GATA1/2 regulation **(Fig. 3I)**. However, despite significant overlap between genes induced by G9a/GLP perturbation and those induced by hydroxyurea (higher than by chance with p = 0.005) **(Fig. S6F)**, there was also a large subset of genes activated only by hydroxyurea, consistent with G9a inhibition and WIZ degradation shifting the global transcriptional profile toward the state induced by hydroxyurea without fully recapitulating the transcriptional response to replication stress **(Fig. S6C)**. This is likely explained by induction of genes related to the replication stress response as well as additional epigenetic changes such as loss of H3K27me3 at alternative lineage loci as described above **(Fig. S5F-L)**.

### G9a is limiting in the soluble cell fraction and remains chromatin bound during replication stress

We next sought to understand why binding of G9a complex members and BCL11A, as well as associated repressive histone marks, are reduced at their target genomic loci upon hydroxyurea treatment. We first compared levels of G9a, WIZ, BCL11A, H3K9me2, and H3K9me3 protein in HUDEP-2 whole cell lysates, the soluble fraction (both nuclear and cytosolic), and the non-soluble (chromatin-bound) fraction before and after treatment with hydroxyurea or aphidicolin for 24 hours. Surprisingly, the total cellular levels of all these proteins were unchanged by drug treatments, and these factors were neither depleted in the chromatin-bound fraction nor increased in the soluble fraction **(Figs. 4A-C)**. Moreover, RNA levels for BCL11A or the G9a/GLP complex members GLP (encoded by *EHMT1*), G9a (encoded by *EHMT2*), WIZ, and ZNF644 also did not change following hydroxyurea treatment **(Fig. S7A)**. Taken together, these data argue that the reduced CUT&RUN peak signal for G9a, WIZ, and BCL11A upon hydroxyurea treatment cannot be explained by decreased expression or reduced total chromatin binding of corepressors. Rather, these factors appear to stay on chromatin but redistribute away from their original genomic binding sites during replication stress.

**Figure 4:**
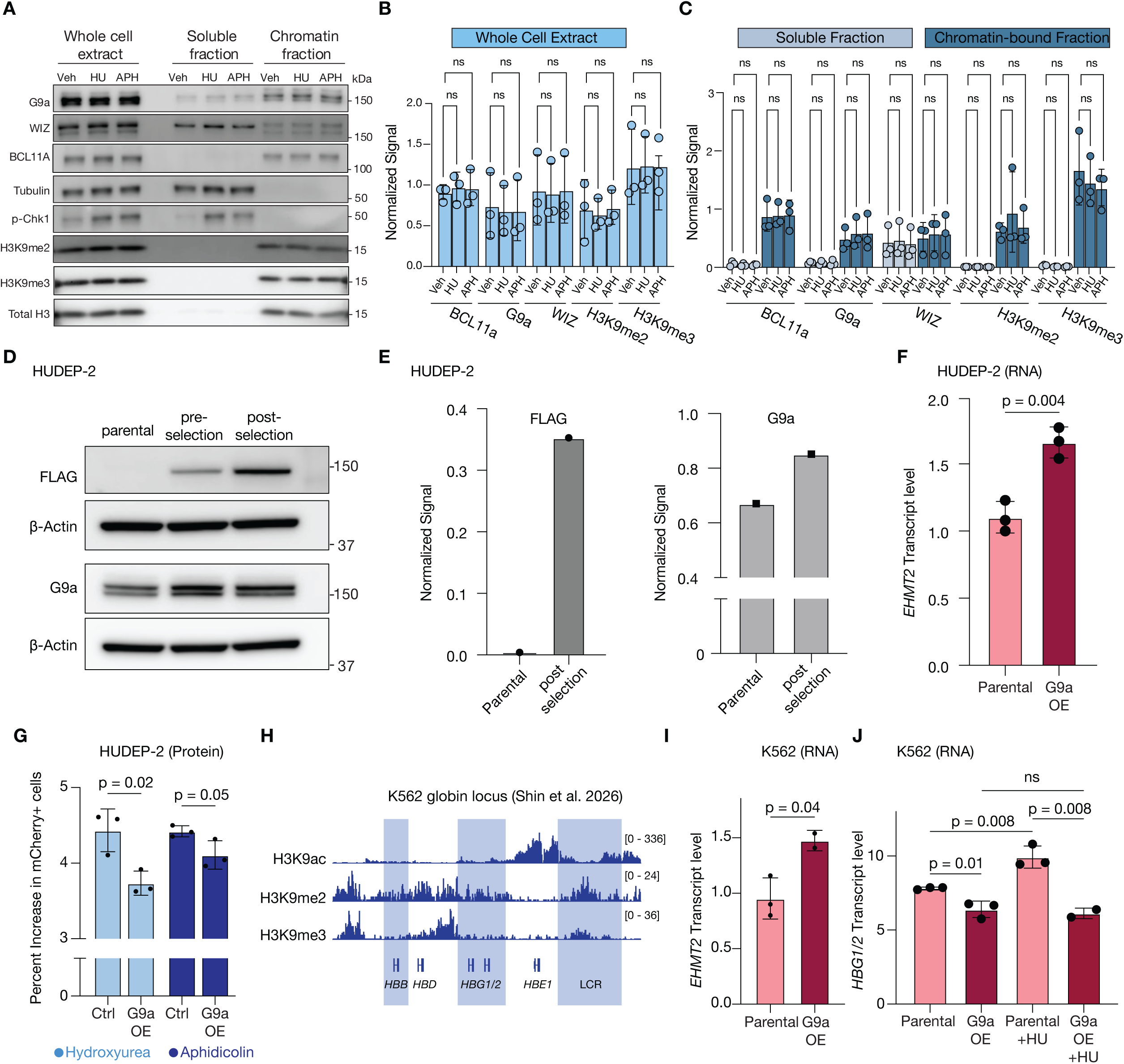
G9a levels are limiting for repression of gene expression. **(A)** Representative immunoblot analysis of G9a, WIZ, BCL11A, H3K9me2 and H3K9me3 in HUDEP-2 whole cell extract, soluble, and chromatin-bound fractions from cells treated with vehicle (Veh), hydroxyurea (HU), or aphidicolin (APH) as indicated. Tubulin serves as a loading control for the soluble fraction and total H3 as a loading control for the chromatin-bound fraction, while Ser345-phosphorylated CHEK1 (p-Chk1) is a marker of DNA replication stress. **(B–C)** Quantification of immunoblot signals from (A) along with other replicates (n = 3, blots not shown) for (B) whole cell extract and (C) soluble and chromatin-bound fractions as indicated. Signal intensities are normalized to indicated loading controls. Statistical significance was determined via two-way ANOVA and p-values > 0.05 are indicated as ns (not significant). **(D)** Representative immunoblot of FLAG and G9a signal in parental HUDEP-2 cells, HUDEP-2 cells transduced with a FLAG-G9a overexpression construct containing a puromycin resistance gene prior to selection, and similarly transduced HUDEP-2 cells selected with puromycin for 72 hr (hereafter referred to as G9a OE cells). β-actin serves as a loading control. **(E)** Quantification of the representative immunoblot shown in (D). FLAG and G9a signal is normalized to their corresponding β-actin loading control. **(F)** *EHMT2* (G9a) RNA transcript levels in parental HUDEP-2 cells and G9a OE HUDEP-2 cells (n=3). p-value was calculated using Student’s t-test. **(G)** Bar plot comparing the increase in the percent of mCherry+ cells following treatment of parental and G9a OE HUDEP-2 cells with either HU or aphidicolin (APH) as indicated (n=3). p-values were calculated using Student’s t-test. **(H)** Genome tracks for H3K9ac, H3K9me2, and H3K9me3 ChIP-seq signal in K562 cells, data derived from previously published work^65^. Highlighted regions denote globin gene clusters and the locus control region (LCR) as indicated. **(I)** *EHMT2* (G9a) RNA transcript levels measured by qRT-PCR in parental K562 cells and K562 G9a OE cells as indicated. qRT-PCR signal is normalized to *PPIB* and *RPLP0*. P-values were calculated using Student’s t-test (n=3). **(J)** *HBG1/2* RNA transcript levels measured by qRT-PCR in parental K562 cells and G9a OE K562 cells with and without HU treatment. qRT-PCR signal is normalized to *PPIB* and *RPLP0.* P-values were calculated using Student’s t-test (n=3).

Levels of G9a were much lower in the soluble fraction relative to the chromatin fraction **(Figs. 4A-C)**, suggesting that there may be insufficient free protein to replace what is lost upon redistribution away from its original genomic binding sites. To test this hypothesis, we overexpressed FLAG-tagged G9a in HUDEP-2 cells **(Figs. 4D-F)** and assessed the degree of HbF induction following treatment with either hydroxyurea or aphidicolin. Strikingly, even slight overexpression of G9a reduced the degree of HbF induction **(Fig. 4D-G)**. We also tested this hypothesis in the K562 erythroleukemia cell line, where H3K9me2, but not H3K9me3, is enriched at *HBG1*, *HBG2*, *HBB*, and the locus control region **(Fig. 4H)**. Overexpression of G9a in these cells completely abrogated the increase in *HBG* expression observed during replication stress **(Figs. 4I-J, S7B-C)**. These data suggest that G9a corepressor levels are limiting for gene repression in our erythroid models, such that redistribution of G9a to other chromatin sites upon replication stress could be involved in the activation of *HBG* and other G9a and H3K9me2-bound genes.

### Replication stress causes accumulation of corepressors and repressive histone marks at replication forks

We hypothesized that G9a is recruited away from its genomic binding sites towards stalled forks during replication stress. Because the genomic sites of fork stalling will vary following hydroxyurea or aphidicolin treatment across cells in different parts of S phase, we did not expect to see accumulation of G9a and H3K9me2 at specific genomic loci within our CUT&RUN data. We therefore used three orthogonal methods to test the hypothesis that G9a moves to sites of replication stress. First, we asked whether replication stress causes replicative DNA to be enriched within the pool of H3K9me2-bound chromatin. To test this, we developed a modified CUT&RUN approach coupled to EdU click chemistry (Click-CUT&RUN) to quantify the enrichment of nascent DNA within H3K9me2-associated chromatin. Briefly, we labeled nascent DNA with the thymidine analog 5-ethynyl-2-deoxyuridine (EdU) in cells treated with either vehicle or hydroxyurea **(Fig. 5A)**, then performed CUT&RUN for H3K9me2. After purifying DNA fragments associated with H3K9me2, we used CuAAC click chemistry^46^ to biotinylate EdU-labeled nascent DNA and quantified the amount of labeled DNA using streptavidin-HRP immunoblotting. Using this approach, we observed increased levels of EdU as a percentage of H3K9me2-labeled DNA upon hydroxyurea treatment compared to vehicle treatment, whereas EdU as a percentage of total DNA was unchanged **(Fig. 5B)**. This observation indicated that replication stress leads to an enrichment of replicative DNA within the total nuclear pool of H3K9me2-bound chromatin, suggesting that factors that deposit H3K9me2 may also be enriched at stalled replication forks.

**Figure 5:**
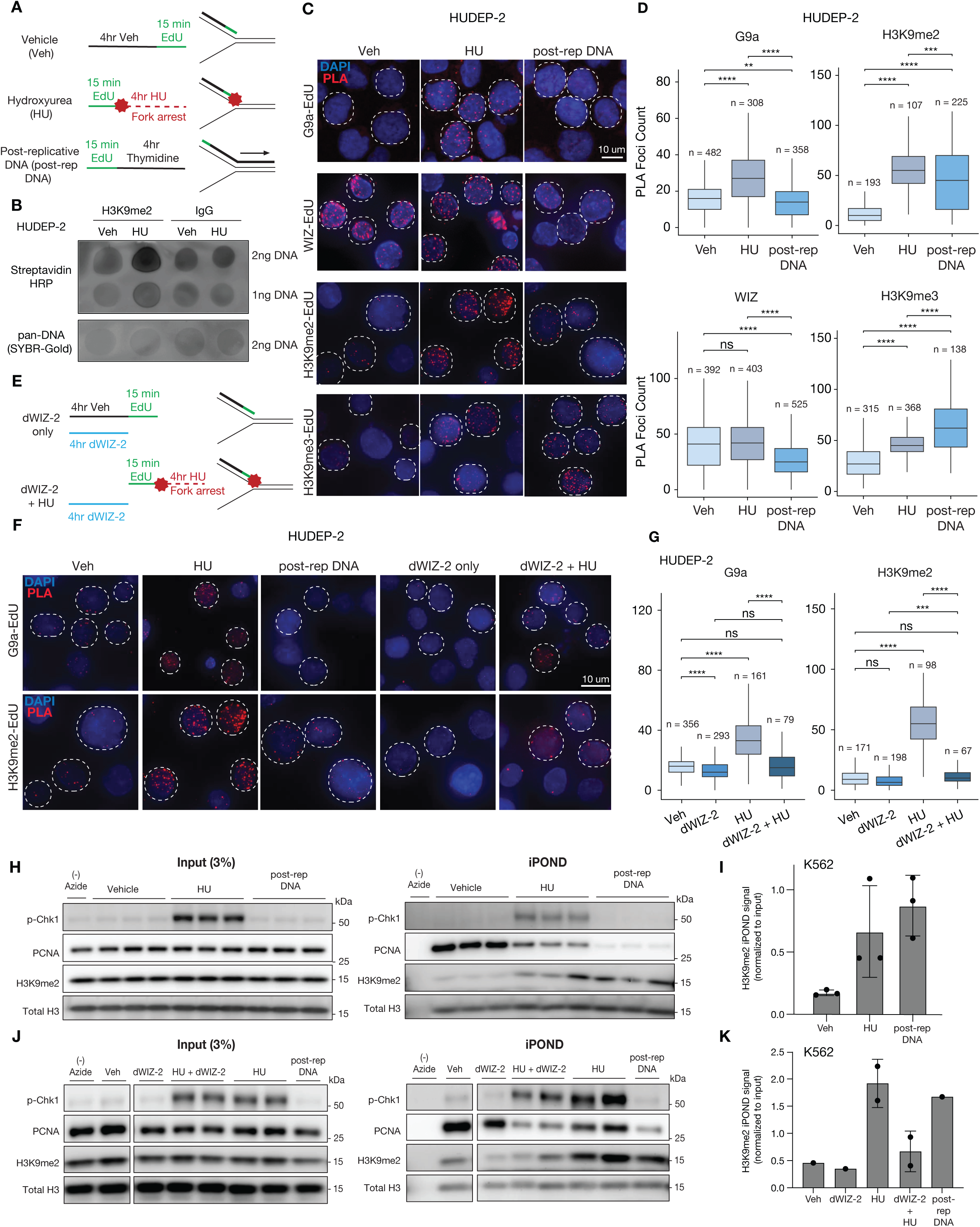
Replication stress promotes redistribution of G9a and H3K9me2 to stalled replication forks. **(A)** Schematic of EdU labeling and compound treatment for a subset of assays shown in this figure. Cells were labeled with EdU under three conditions: Vehicle (Veh), with 4 hr vehicle treatment followed by 15 min EdU pulse, Hydroxyurea (HU), with 15 min EdU pulse followed by 4 hr hydroxyurea (HU) treatment at a dose that fully arrests fork progression, and Post replication DNA (post-rep DNA), with a 15 min EdU pulse followed by 4 hr thymidine chase. Timelines of treatment are shown (left) as well as position of EdU labeling indicated in green on replication forks (right). **(B)** Representative streptavidin-HRP immunoblots for EdU-labeled DNA pulled down by H3K9me2 or IgG click-CUT&RUN in HUDEP-2 cells treated with Veh or HU as described in (A). The indicated amount of DNA was loaded in each lane. Equal DNA loading was assessed using a pan DNA stain (SYBR-Gold). **(C)** Representative immunofluorescence images of foci per nucleus for G9a, WIZ, H3K9me2, or H3K9me3 in proximity to nascent DNA (labeled by EdU) using a proximity ligation assay (PLA). PLA signals (red) are shown for each condition described in (A) as indicated, and nuclei are counterstained with DAPI (blue). Cells in S phase, determined by the presence of EdU-labeled DNA, are represented by white dashed circles. Representative selection of EdU-positive and EdU-negative signal is shown in Figure S8A. **(D)** Quantification of PLA foci per nucleus for G9a, WIZ, H3K9me2, and H3K9me3 from the PLA shown in (C). Data are presented as box-and-whisker plots showing median and interquartile range. Statistical significance was calculated using a Wilcoxon test with Benjamini-Hochberg p-value correction (**p < 0.01, ***p < 0.001, ****p < 0.0001; ns, not significant). The number of nuclei assessed for each condition is indicated. **(E)** Schematic of EdU labeling and dWIZ-2/HU treatment for subset of assays shown in this figure. Cells were labeled with EdU and treated with dWIZ-2 under two conditions: dWIZ-2 only, with 4 hr dWIZ-2 treatment followed by 15 min EdU pulse, and dWIZ-2 + HU, with 4 hr dWIZ-2 treatment followed by a 15 min EdU pulse followed by 4 hr HU. Timelines of treatment are shown with dWIZ-2 shown in blue (left) as well as position of EdU labeling indicated in green on replication forks (right). **(F)** Representative immunofluorescence images of PLA signals (red) for G9a or H3K9me2 in proximity to EdU for vehicle-treated, HU, post-rep DNA, dWIZ-2 only, and dWIZ-2 + HU conditions as described in (A) and (E). Nuclei are counterstained with DAPI (blue). Cells in S phase are represented by white dashed circles. **(G)** Quantification of PLA foci for G9a and H3K9me2 in proximity to EdU in cells treated with vehicle or HU with or without degradation of WIZ through dWIZ-2 pretreatment as shown in (E). Data are presented as box-and-whisker plots. Statistical significance was calculated using a Wilcoxon test with Benjamini-Hochberg p-value correction (**p < 0.01, ***p < 0.001, ****p < 0.0001; ns, not significant). The number of nuclei assessed for each condition is indicated. **(H)** Representative immunoblot of 3% pulldown input (left) and proteins associated with nascent DNA identified by iPOND (isolation of proteins on nascent DNA)^66^ (right) in K562 cells treated with vehicle or HU, or following a post-replicative chase (see (A)). In iPOND, EdU incorporated into nascent DNA is conjugated to biotin azide by click chemistry^60^, enabling affinity purification of nascent DNA and associated proteins. The (-) azide sample, in which biotin azide was omitted from the click reaction, serves as a negative control for nonspecific protein recovery during purification. **(I)** Quantification of H3K9me2 iPOND signal from (H), normalized to H3 iPOND signal and the input H3K9me2/H3 ratio, at vehicle and HU-treated nascent DNA and at post-replication DNA (n=3). **(J)** Representative immunoblot of 3% pulldown input (left) and iPOND fractions (right) from Veh, HU, post-rep DNA, dWIZ-2 only, and dWIZ-2 + HU (see (A) and (E)) from K562 cells. **(K)** Quantification of H3K9me2 iPOND signal from (J), normalized to H3 iPOND signal and the input H3K9me2/H3 ratio.

Second, we used a proximity ligation assay (PLA)^47^ to quantify interactions of stalled forks with G9a or repressive histone marks. We again labeled nascent DNA by pulsing cells with EdU for 15 minutes, then treated cells with high doses of hydroxyurea to initiate fork stalling **(Fig. 5A)**. As a control, we assessed interactions between repressive machinery and post-replicative DNA by exposing cells to the same 15-minute EdU pulse followed by a thymidine chase for 4 hours **(Fig. 5A)**. Interactions between EdU-labeled DNA and antibody-labeled factors within the 40-nanometer radius detected by PLA appear as foci that can be quantified^47^. We confirmed that PCNA, a replication fork-associated protein, was enriched at both vehicle-treated and hydroxyurea-treated nascent DNA compared to the post-replicative control, and that phospho-Ser33 RPA2 (p-RPA), which is deposited at single-stranded DNA and requires ATR-mediated phosphorylation, was enriched only in hydroxyurea-treated nascent DNA compared to untreated nascent DNA or post-replicative DNA **(Figs. S8B-C)**. Consistent with prior observations^48,49^, we found that the hydroxyurea-treated cells displayed increased colocalization between nascent DNA and H3K9me2. H3K9me3 was also enriched at replication forks in hydroxyurea-treated cells compared to vehicle-treated replication forks **(Figs. 5C-D)**, but it was even more enriched in post-replicative DNA, likely reflecting a mechanism of time-dependent deposition of heterochromatin-associated marks^50^ that is distinct from H3K9me2. In line with the observed accumulation of H3K9me2 at stalled replication forks, we also observed colocalization of G9a and EdU-labeled replication forks during hydroxyurea treatment, and not in vehicle treatment or at post-replicative DNA (**Figs. 5C-D)**. Interestingly, colocalization between WIZ and nascent DNA was high with or without replication stress. This is consistent with both the relatively high levels of WIZ observed in the soluble cell fraction compared to G9a **(Figs. 4A, C)** as well as the large number of WIZ peaks relative to G9a peaks observed via CUT&RUN **(Fig. 2F)** and suggests that unlike G9a, WIZ may not be limiting for chromatin binding. We repeated these experiments in a second HUDEP-2 clone with higher baseline HbF and observed similar results **(Fig. S8D)**. These findings support a model in which replication stress promotes accumulation of G9a at stalled replication forks to deposit repressive chromatin modifications including H3K9me2.

The G9a complex member WIZ, which contains a DNA-binding domain, has been hypothesized to play a role in recruiting G9a to its functional sites on chromatin^20^. Thus, we sought to test whether the presence of WIZ at replication forks was necessary for the recruitment of G9a and subsequent deposition of H3K9me2 at stalled replication forks. We performed PLA after treating cells with the WIZ-degrading compound dWIZ-2 for four hours prior to EdU pulse and hydroxyurea treatment (**Fig. 5E**). Even in unstressed conditions, dWIZ-2 treatment led to a decrease in colocalization of WIZ and G9a with replication forks (**Figs. 5F, S8E**). Upon hydroxyurea treatment, addition of dWIZ-2 completely abolished the increased colocalization between replication forks and both G9a and H3K9me2 (**Figs. 5F-G**). These data suggest that WIZ is required for the recruitment of G9a, and subsequent deposition of H3K9me2, at stalled replication forks.

As a third method for testing whether G9a and H3K9me2 accumulate at stalled forks, and to understand whether accumulation of repressive epigenetic marks at stalled replication forks was observed in other cell types, we performed isolation of proteins on nascent DNA (iPOND)^51,52^ in K562 cells^53^, where G9a overexpression had also abrogated *HBG* induction in response to hydroxyurea treatment **(Fig. 4J)**. In brief, we labeled nascent DNA for 15 minutes followed by stalling replication forks with hydroxyurea for one hour before formaldehyde crosslinking, biotinylation of EdU, purification of biotinylated DNA-protein complexes, and immunoblotting of enriched proteins. We observed robust increases in H3K9me2 at stalled replication forks relative to normal replication forks (**Figs. 5H-I**). Accumulation of H3K9me2 was suppressed by pretreatment of cells with dWIZ-2 prior to fork stalling (**Figs. 5J-K**). These data confirm that stalled replication forks accumulate H3K9me2 in a WIZ-dependent manner in a separate cell line. Together with the PLA results, these data support the hypothesis that during replication stress, G9a is recruited from its target genomic loci to stalled replication forks where it promotes deposition of H3K9me2^49^. Genes associated with reduced G9a binding and H3K9me2 deposition are then activated, providing a mechanism by which replication stress activates expression of a subset of primed genes.

## Discussion

Here we show that replication stress activates epigenetically primed genes by redistributing euchromatin-specific corepressors from their canonical binding sites to stalled replication forks. In erythroid progenitor cells, *HBG* is an example of a primed locus that relies on the euchromatic histone methyltransferase G9a to maintain H3K9me2-mediated gene silencing. We also observe epigenetic and transcriptional derepression of GATA1/2-associated loci important for erythroid differentiation. Using multiple orthogonal methods, we demonstrate that G9a is physically recruited to stalled replication forks in a WIZ-dependent manner, and because G9a levels are limiting in cells, it is depleted from its canonical target sites, leading to gene activation.

These data suggest an explanation for how hydroxyurea reactivates HbF expression. In the 1980s, it was postulated that perturbations such as stress erythropoiesis or replication inhibitors selectively amplify a subset of erythroid progenitors, and that these cells are either intrinsically biased for HbF expression or have mechanisms for maintaining a HbF-expressing state. Our data indicate that erythroid progenitors can transition flexibly between A and F states, and that short-term epigenetic factors persisting at least through several cell divisions influence the degree of HbF induction upon replication stress. Of note, F cells sorted from a random population, or following pretreatment with hydroxyurea, gradually revert to the A state unless replication stress is sustained. This suggests that the F cell state acquired during replication stress requires active maintenance, potentially through continued G9a/GLP sequestration at stalled DNA replication forks. In combination with recent data suggesting that there are few transcriptional differences between F and A progenitors^28^, we favor a more general model where some erythroid progenitors are, by chance, more likely to express HbF upon replication stress, possibly as a result of single-cell variation in corepressor availability.

Our findings extend recent observations that have linked replication stress to chromatin remodeling at stalled replication forks. For example, stalled replication forks also accumulate chromatin activators such as PTIP^54^ and corepressors such as SETDB1 and POGZ^55^. If levels of these factors are also limiting in cells, we hypothesize that, like G9a, their redistribution to stalled forks could deplete them from target sites across the genome. Indeed, replication stress has been shown to induce lineage-specific changes in gene expression and cell state across multiple contexts, an observation that is difficult to explain solely through activation of a replication stress or DNA damage signaling pathway^4^. Redistribution of corepressors could explain why primed genes, which are unique to the cellular context, are particularly susceptible to derepression following replication stress and explain how replication stress activates a lineage-specific change in gene expression state.

How stalled replication forks recruit the G9a complex, as well as whether all the complex members are recruited together or if WIZ is already present, is not known. As WIZ is directly required for recruitment of G9a, it may be that there are active recruitment mechanisms, rather than passive retention of factors on nascent chromatin. Alternatively, the partitioning of actively replicating forks into “replication factories” could enable passive recruitment of corepressors through concentration of low-affinity interactions. Interestingly, several published iPOND-MS and NCC-SILAC-MS datasets^56,57^ comparing replication fork-associated proteomes in untreated versus hydroxyurea-treated cells have found depletion rather than enrichment of the G9a/GLP complex, which is inconsistent with our PLA data as well as our iPOND data showing enrichment of H3K9me2 deposition. We hypothesize that this may be due to the differences in the biochemical interactions captured by different approaches. Lastly, our data suggest that the ATR/Chk1-mediated replication stress response is not necessary for G9a/WIZ recruitment to stalled forks, consistent with prior work in leukemia models suggesting that inhibition of ATR does not suppress the cell state changes triggered by replication stress. The contribution of other signaling pathways activated during replication stress remains unclear^58^. Future work will need to define the full complement of regulatory factors that are recruited to stalled forks as well as the mechanisms that mediate their recruitment.

Our work provides additional evidence that replication stress contributes to broader transcriptional and epigenetic effects than previously understood. In pathological states such as cancer, both endogenous replication stress, caused by rapid proliferation and nutrient limitation, and on-target replication defects driven by chemotherapy and targeted therapies such as PARP inhibitors could conceivably provide surviving cells with the epigenetic plasticity to transition into new cell states including those that confer drug resistance or metastatic capability. Individuals with germline defects in DNA replication or repair, such as Fanconi anemia, exhibit bone marrow failure, increased cancer predisposition, and endocrine deficiencies, raising the possibility that replication stress may contribute to the generation of cell states that drive these phenotypes. Lastly, as primed chromatin states are critical mechanisms to promote lineage bias during organismal development, replication stress-induced redistribution of chromatin corepressors may represent a broader mechanism linking DNA replication dynamics to cell fate transitions in normal development as well as in disease.

## Acknowledgements

We thank members of the Vander Heiden laboratory as well as Yadira Soto-Feliciano and members of her lab for helpful discussions. We also thank the Koch Institute’s Robert A. Swanson (1969) Biotechnology Center for technical support, especially Jeffrey Kuhn and Jeffrey Wyckoff in the Spatial Imaging Core, Glenn Paradis, Michael Tramontanis, Michele Griffin, and Ella Skinner-Sloan in the Flow Cytometry Core, and Stuart Levine in the MIT BioMicro Center.

## Funding

G.E.O acknowledges support from the MIT Graduate Dean of Science Fellowship. B.T.D. acknowledges support from the National Heart, Lung, and Blood Institute (F30HL156404). D.E.B. acknowledges support from the National Heart, Lung, and Blood Institute (R01HL167513) and the St. Jude Sickle Cell Disease Collaborative Research Consortium. M.G.V.H. acknowledges support from the National Cancer Institute (R35CA242379, P30CA14051), the MIT Center for Precision Cancer Medicine, and the Ludwig Center at MIT.

## Author Contributions

B.T.D., G.E.O., and M.G.V.H. conceived of the project; G.E.O., B.T.D., E.M.J., A.P.W., C.M.C., and S.A.N. conducted most experiments; P.J.C. conducted RNA-seq analysis. D.E.B. provided critical reagents and advice. B.T.D. and G.E.O. wrote the paper, with input from M.G.V.H. and all co-authors. M.G.V.H. obtained funding.

## Competing Interests

M.G.V.H. discloses that he is a scientific advisor for Agios Pharmaceuticals, Pretzel Therapeutics, Lime Therapeutics, Faeth Therapeutics, Droia Ventures, Verdandi Therapeutics, Alterris Oncology, and Auron Therapeutics. M.G.V.H. also discloses that he is on the editorial board of Molecular Cell. All remaining authors declare no competing interests.

## Lead Contact

Further information and requests for resources and reagents should be directed to and will be fulfilled by Matthew G. Vander Heiden.

## Methods

### Cell Culture

HUDEP-2 (ATCC) human erythroid progenitor cells were cultured in SFEM II media (Stem Cell Technologies, 09665) supplemented with 3U/mL erythropoietin (Stem Cell Technologies, 78007.1), 50ng/mL stem cell factor (PeproTech, 300-07), 2µM dexamethasone (Sigma-Aldrich, D4902), and 1µg/mL doxycycline (Sigma-Aldrich, D9891). Doxycycline was added fresh to culture media every 2-3 days. Cell density was kept between 1 × 10⁵ and 8 × 10⁵ cells/mL. Primary human CD34⁺ hematopoietic stem and progenitor cells were purchased from StemCell Technologies (StemCell Technologies, 70002.1), who obtained them from healthy, deidentified donors. They were cultured under CD34^+^ expansion conditions (SFEM II media (StemCell Technologies, 09655) with Erythroid Expansion Supplement (StemCell Technologies, 02692)) for 7 days. K562 human chronic myelogenous leukemia cells (ATCC, CCL-243) were cultured in RPMI 1640 media (Corning, 15-040-CV) supplemented with 10% fetal bovine serum (Gibco, A52567-01) and 4mM L-glutamine (Cytiva, SH30034.01). Cell density was kept between 1 × 10⁵ and 1 × 10⁶ cells/mL. All cells were maintained at 37 °C with 5% CO₂, passaged every 2-3 days, and routinely tested for mycoplasma.

### Genetic Modification of Cell Lines

For all lentiviral transductions, lentiviral particles were generated by transfecting 293T cells (ATCC, CRL-3216) with a 4:2:1:1 ratio of the TARGET : pMDLg : pMD2.G : pRSV-REV plasmids using Transit 293T reagent (Mirus Bio, MIR 2700). After 48 hours, supernatant was clarified through a 0.45μm PES filter and either stored at −80°C or added fresh to target cells. All cells were transduced using 250μl viral supernatant and 250μl cells at a concentration of 3 x 10^5^ cells/mL with polybrene (EMD Millipore, TR-1003-G) added at a final concentration of 10μg/mL. *HBG1/2 reporter cell line:* A fluorescent transcriptional reporter for *HBG2* was expressed as previously described^27^. Briefly, CRISPR/Cas9-mediated genome editing was used to introduce a reporter cassette downstream of the endogenous gamma-globin promoter. Correct integration was confirmed by PCR and sequencing and reporter activity assessed by flow cytometry. Clonal lines with induction of fluorescence upon HU treatment were selected for downstream experiments. The primers used for validation are listed in Table S1.

#### G9a Overexpression

A G9a overexpression construct was created by combining the G9a coding sequence from pLenti6-MK1-EHMT2-V5 (Addgene, #31113) with a pLJM1 backbone (Addgene, #19319) conjugated to a flexible linker and 3xFLAG-tag sequence (AGGADYKDHDGDYKDHDIDYKDDDDK) (Addgene #262548). HUDEP-2 cells and K562 cells were transduced as described above and selected with 1µg/mL puromycin (Sigma-Aldrich, P8833) for 72 hours. Cells were passaged outside of selection media for a minimum of 48 hours before experimental use.

#### HRI knockdown

HRI knockdown was performed using shRNA constructs (TRCN0000010229 and TRCN0000010231 from the Broad Genome Perturbation Platform) expressed from a pLKO.1 backbone (Addgene #8453). HUDEP-2 cells were transduced as described above and selected with 1µg/mL puromycin.

### Chemical compounds

The following drugs were used to induce replication stress: hydroxyurea (HU; Sigma-Aldrich, H8627 and EMD Millipore, 400046), aphidicolin (APH; Sigma-Aldrich, A0781), etoposide (Sigma-Aldrich, E1383), decitabine (DEC; MedChemExpress, HY-A0004), methotrexate (Sigma-Aldrich, A6770), camptothecin (Santa Cruz Biotechnology, SC-200871), gemcitabine (GEM; MedChemExpress, HY-17026), pomalidomide (POM; TCI Chemicals, P2074), and cytarabine (Selleck Chemicals, S1648). ATR signaling was inhibited using AZ20 (Selleck Chemicals, S7050 and MedChemExpress, HY-15557). To disrupt the G9a/GLP complex, cells were treated with dWIZ-2 (Selleck Chemicals, E4661) and UNC-0646 (Cayman Chemical Company, 11085). Efficiency of ATR inhibition and WIZ degradation was confirmed by immunoblotting. All drugs were stored at a stock concentration of 10mM in DMSO (Sigma-Aldrich, D2650) at −20°C except for hydroxyurea, which was stored at a stock concentration of 500mM in water at −20°C, dWIZ-2, which was stored at a stock concentration of 20mM in DMSO at −20°C, and UNC-0646, which was stored at a stock concentration of 1mM in DMSO at −20°C.

### Flow Cytometry Analysis and Sorting

Flow cytometry analysis was performed on a BD FACSymphony A3 flow cytometer. For cell viability analysis, cells were incubated for 15 minutes with 1µg/mL DAPI (Thermo Fisher Scientific, D1306) in PBS. For surface staining, the following antibodies were used at a 1:100 dilution: CD71-SuperBright436 (Invitrogen, clone OKT9, 62-0719-42) and CD235a-APC (BD

Biosciences, clone GA-R2, 561775). Cells were washed once with PBS, then stained for 15 minutes on ice followed by a single wash with ice-cold PBS and resuspension in PBS with 1µg/mL DAPI (Thermo Fisher Scientific, D1306) for live/dead cell discrimination. For intracellular flow cytometry, the following primary antibodies were used at a 1:100 dilution: p-Chk1 (Cell Signaling Technologies, 2341), HbF-FITC (Novus Biologicals, NB110-41084F), and HbF-APC (Thermo Fisher Scientific, MHFH05). The following secondary antibodies were used at 1:1000 dilution: AF488-conjugated anti-mouse highly cross-adsorbed secondary antibody (Invitrogen, A11029), and AF647-conjugated anti-rabbit highly cross-adsorbed secondary antibody (Invitrogen, A21244). 100,000 cells were pelleted and washed once with PBS. Cells were then fixed in 4% paraformaldehyde (Electron Microscopy Sciences, 15710) (made fresh) for 15 minutes at room temperature and permeabilized in 0.2% Triton-X-100 (Thermo Fisher Scientific, 327371000) for 20 minutes at room temperature. Cells were then incubated with primary antibodies for 30 minutes at 4°C protected from light. Cells were washed once with ice-cold FACS buffer (1% FBS (Gibco, A52567-01) in PBS). If required, cells were then incubated with secondary antibodies for 30 minutes at 4°C protected from light. Cells were then washed with FACS buffer and moved to flow tubes for data analysis. Staining for phosphorylated CHEK1 was performed with Tris-buffered saline (TBS) instead of PBS to prevent phosphate-antibody interactions. Sorting was performed on BD FACSAria machines at the Koch Institute flow cytometry core, using a 70μm nozzle. All cells were sorted into fresh cell culture media with 1% Pen/Strep (Corning, 30-002-CI), which was removed after 48 hours. Gating and downstream analysis were performed using FACSDiva (BD) or FlowJo v10 software.

### CUT&RUN

CUT&RUN was performed using the Epicypher CUTANA CUT&RUN kit (Epicypher, 14-1048), using between 100,000 and 500,000 cells per sample for each experiment. Cells were counted twice and the result was averaged, and 5% of this number of freshly thawed mouse ER-Hoxa9 myeloid progenitors^59^ were added as a spike-in. Cells were then immobilized on concanavalin A-coated beads, permeabilized, and incubated overnight at 4°C with 0.5µg of antibodies against G9a (Cell Signaling Technologies, 68851), WIZ (Novus, NBP1-80586), BCL11A (Cell Signaling Technologies, 75432), H3K9me2 (EMD Millipore, 05-1249), H3K9me3 (Abcam, AB176916), H3K9ac (Cell Signaling Technologies, 9649), H3K27ac (EMD Millipore, MABE647), H3K27me3 (Cell Signaling Technologies, 9733), H3K4me1 (EMD Millipore, 07-436), H3K4me3 (EpiCypher, 13-0028), or IgG (EpiCypher, 13-0042k). Protein A/G–micrococcal nuclease was used to cleave antibody-bound chromatin and released DNA fragments were purified and size-selected via SPRI magnetic beads (Sergi Lab Supplies, #1040), using a 1:1 bead to DNA ratio for histone libraries and a 1.6:1 bead to DNA ratio for transcription factor libraries in order to capture smaller fragments. Sequencing libraries were prepared with the NEBNext® Ultra™ II DNA Library Prep Kit for Illumina (New England Biolabs, E7645L) according to the manufacturer’s protocol, except that for transcription factor libraries, end prep was performed at 50°C instead of 65°C due to shorter fragment length. Libraries were multiplexed using custom-designed combinatorial dual index primers (36 i7 and 8 i5 primers) (Table S2). Sequencing was performed on an Illumina NovaSeq X Plus at the University of California, Irvine, Genomics Research and Technology Hub, targeting an average of 20 million reads per sample.

For data analysis, FASTQ files were demultiplexed and mapped first to the mm10 genome to quantify and remove spike-in reads, then to the hg38 genome using Bowtie2 v2.4.2 (with parameters --dovetail -I 10 -X 700 --very-sensitive-local --local). Reads corresponding to blacklisted regions were removed, and tracks were normalized using the bamCoverage tool from deepTools (v3.5.0) (with parameters -bs 1 -e 200). Duplicate reads (inferred using Picard) comprised less than 15% of total reads in all samples and were thus not removed. Peaks were called using MACS2 (with parameters -q 0.01 --nomodel) and filtered to retain only those that appeared in all biological replicates of a given sample.

### Cellular Fractionation

Approximately 1 million HUDEP cells treated with 500µM HU or 2.5µM APH or left untreated for 24 hours were split evenly into a whole cell extract (WCE) and extraction tube and pelleted. The supernatant was discarded and the cell pellets were washed twice with ice-cold PBS. The pellets in the WCE tubes were resuspended in 150µL CSK Buffer (10mM PIPES pH 7.0 (Boston BioProducts, BB-2284), 300mM sucrose (Sigma-Aldrich, S9378), 3mM MgCl_2_ (Thermo Fisher Scientific, AM9530G), 1mM EGTA (Fisher Scientific, NC9339264), 0.5% Triton-X-100 (Thermo Scientific, 327371000), 1mM DTT (Sigma-Aldrich, 646563-10X), supplemented with Phosphatase/Protease Inhibitor cocktail (Thermo Scientific, 78440)) with 50U/mL benzonase (EMD Millipore, 70746-3) and rotated at 4°C for 30 minutes. The pellets in the extraction tubes were resuspended in 170µL CSK buffer after which they were placed on ice for exactly 10 minutes, and subsequently spun down at 1,500 x g for 3 minutes at 4°C. The supernatant of the extraction tubes was transferred to clean tubes and spun down at maximum speed for 10 minutes at 4°C. Following centrifugation, 150µL of the supernatant was transferred to a clean tube as the soluble fraction. The remaining chromatin pellet was washed twice with 1mL of CSK buffer and the chromatin pellet was resuspended in 150µL CSK buffer with 50U/mL benzonase and rotated at 4°C for 30 minutes. All samples (WCE, soluble, and chromatin) were then normalized using the Pierce BCA Protein Assay Kit (Thermo Scientific; #23225) with BSA as a standard. 50µL of 4X LDS (GenScript, M00676) with 10% beta-mercaptoethanol (Sigma-Aldrich, M6250) was added to each sample, followed by incubation at 95°C for 10 minutes. All fractions were then analyzed via immunoblot.

### Immunoblotting

The following primary antibodies were used, all at a dilution of 1:1000: G9a (Cell Signaling Technologies, 68851), WIZ (Novus, NBP1-80586), BCL11A (Cell Signaling Technologies, 75432), H3K9me2 (EMD Millipore, 05-1249), H3K9me3 (Cell Signaling Technologies, 9733), p-S345 Chk1 (Cell Signaling Technologies, 2341), p-T68 Chk2 (Cell Signaling Technologies, 2197), Chk1 (Cell Signaling Technologies, 37010), Chk2 (Cell Signaling Technologies, 6334), beta-actin (Cell Signaling Technologies, 8457), beta-tubulin (Cell Signaling Technologies, 2146), vinculin (Cell Signaling Technologies, 13901). The following secondary antibodies were used: HRP-linked anti-rabbit IgG (Cell Signaling Technologies, 7074; used at 1:5000) and HRP-linked anti-mouse IgG (Cell Signaling Technologies, 7076; used at 1:2000). Briefly, 500,000 cells were washed twice with 1mL ice-cold PBS, pelleted, lysed using RIPA buffer (Boston BioProducts, BP-115) at a concentration of 5 million cells/mL, and then centrifuged at maximum speed to pellet cell debris. Samples were normalized using the Pierce protein assay kit (Thermo Scientific, 23225) using bovine serum albumin (GoldBio, A-420-500) in water as a standard. The supernatant was then combined in a 3:1 ratio with 4X LDS (GenScript, M00676) with 10% beta-mercaptoethanol (Sigma-Aldrich; M6250) and the lysate was heated at 70°C for 10 minutes. Lysates were resolved by SDS-PAGE using Bolt 4-12%, Bis-Tris Plus WedgeWell polyacrylamide gels (Invitrogen by Thermo Fisher Scientific, NW04120BOX) and proteins were transferred onto 0.2µm PVDF membranes (EMD Millipore, ISEQ00010) using the Bio-Rad Mini Trans-Blot Electrophoretic Transfer Cell (Bio-Rad, 1703930) at 100V for 90 minutes. Membranes were incubated with primary antibodies overnight at 4°C and incubated with secondary antibodies for 30 minutes at room temperature at concentrations described above, then imaged using Western Lightning Plus ECL Oxidizing Reagent Plus and Western Lightning Plus ECL (Revvity, 0RT2655 and 0RT2755) on an Amersham ImageQuant 800 by Cytiva.

### Click Chemistry detection of EdU

In order to perform CuAAC click chemistry^60^, the protocol for the Click-iT Plus EdU Flow Cytometry Assay Kit for Alexa Fluor 647 and Alexa Fluor 488 (Thermo Fisher Scientific, C10634 and C10633) was used with custom reagents and minor modifications to the manufacturer’s instructions. Briefly, 100,000 cells were pulsed with 10μM 5-ethynyl-2′-deoxyuridine (EdU; Sigma-Aldrich, 900584) for 15 minutes prior to cell collection. After fixation for 15 minutes at room temperature in 100µL of 4% paraformaldehyde (Electron Microscopy Sciences, 15710) (made fresh), permeabilization was performed for 15 minutes with 100μl of 0.2% Triton-X-100 (Thermo Scientific, 327371000). A master mix was made with the following amounts of reagent per sample (added in order): 87μl PBS, 2μl of 100mM Copper (II) Sulfate (Sigma-Aldrich, C1297), 1μl of 500μM picolyl azide (either fluorescent AF488 picolyl azide (Invitrogen by Thermo Fisher, C10641A) or biotin picolyl azide (Sigma-Aldrich, 900912)), and 10µL of 100mM sodium ascorbate (Sigma-Aldrich, A7631). 100μl of this master mix was then added to each sample of permeabilized cells, and the click reaction was allowed to proceed at room temperature for 30 minutes in the dark, then washed once with ice-cold PBS, before proceeding to further staining or downstream analysis.

### Click-CUT&RUN for detection of CUT&RUN signal at nascent DNA

Cells were pulsed for 15 minutes in normal culture conditions with 10μM 5-ethynyl-2′-deoxyuridine (EdU; Sigma-Aldrich, 900584). After 15 minutes, cells were either rapidly washed with excess volumes of CUT&RUN Wash Buffer (19.22mM HEPES pH 7.5 (Sigma-Aldrich, H4034), 144.16mM NaCl (Sigma-Aldrich, S3014), 0.48mM Spermidine (EpiCypher, 21-1005), 1X protease inhibitor tablet (Roche Diagnostics, 82336300) pre-dissolved in 2mL water to make a 25X stock) or dosed with 1mM hydroxyurea (EMD Millipore, 400046) for 4 hours in normal culture conditions to arrest replication forks prior to the wash step. CUT&RUN was then performed using the Epicypher CUTANA CUT&RUN kit (EpiCypher, 14-1048) as described above. 10ng of purified DNA was then conjugated to biotin picolyl azide (Sigma-Aldrich, 900912) using CuAAC click chemistry^60^ as described previously, followed by repurification with SPRI magnetic beads (Sergi Lab Supplies, #1040) and normalization of DNA concentration across samples. Log2 dilution series were then prepared using 1X saline-sodium citrate buffer (SSC; Invitrogen by Thermo Fisher Scientific, AM9763), starting from 2ng/µL of DNA. 1µL of DNA was dotted onto a damp 0.2μm nitrocellulose membrane (Bio-Rad, 1620112) that was pre-equilibrated in 1X SSC for 5 minutes. Samples were loaded slowly to limit spreading. DNA was then fixed onto the membrane by baking at 80°C for 2 hours. The membrane was blocked in 5% BSA (GoldBio, A-420-500) in TBST for 1 hour at room temperature, followed by a 45-minute incubation with streptavidin-HRP (Thermo Scientific, #N100; 1:10000 in 1% BSA in TBST). The membrane was then washed three times for ten minutes in TBST and imaged using Western Lightning Plus ECL Oxidizing Reagent Plus and Western Lightning Plus ECL (Revvity, 0RT2655 and 0RT2755) on an Amersham ImageQuant 800 by Cytiva. As a pan DNA stain loading control, following chemiluminescent imaging, the membrane was rinsed briefly with TBST, incubated with SYBR Gold (Invitrogen by Thermo Fisher, #S11494; 1:10000 in TBS) for 10 minutes at room temperature, rinsed briefly with TBS and imaged using fluorescence imaging of the Cy3 channel.

### Proximity Ligation Assay (PLA)

Proximity ligation assays (PLA) were performed using the Duolink® In-Situ PLA system according to the manufacturer’s instructions. Nascent DNA was labeled with 10μM 5-ethynyl-2′-deoxyuridine (EdU; Sigma-Aldrich, 900584) for 15 minutes immediately prior to fixation or fork stalling with 1mM hydroxyurea (EMD Millipore, 400046) for four hours. Where indicated, cells were treated with 20μM dWIZ-2 (Selleck Chemicals, E4661) for 4 hours prior to EdU labeling. For post-replicative DNA measurements, cells were pulse-labeled with 10μM EdU for 15 minutes at 37 °C, washed with fresh media containing 10μM thymidine (Sigma-Aldrich, 89270), and subsequently incubated in EdU-free medium for 4 hours before fixation. Cells were collected, washed with PBS, and fixed in 4% paraformaldehyde (Electron Microscopy Sciences, 15710) at room temperature for 15 minutes and then permeabilized with 0.2% Triton-X-100 (Thermo Scientific, 327371000) for 15 minutes at room temperature. For PLA measurements involving EdU-labeled DNA, incorporated EdU was conjugated to a biotin picolyl azide (Sigma-Aldrich, 900912) by CuAAC click chemistry^60^ with the click reactions performed as described previously. Subsequently, cells were centrifuged onto poly-D-lysine (Thermo Scientific, A3890401) coated glass slides (Mercedes Medical, MER 7255/90/BL). Slides were then blocked using Duolink Blocking Solution (included in Duolink® in Situ Detection Reagents Red, Sigma-Aldrich, DUO92008) for 60 min at 37°C in a humidified chamber. Primary antibodies were diluted in Duolink Antibody Diluent (included in Duolink® in Situ Detection Reagents Red, Sigma-Aldrich, DUO92008) and incubated with samples overnight at 4°C. To detect proteins or histone modifications in proximity to nascent DNA, samples were incubated with a mouse antibody recognizing biotin (Jackson Immunoresearch Laboratories, 200-02-211) together with a rabbit antibody against the protein or histone modification of interest. Rabbit antibodies used were as follows: G9a/EHMT2 (Cell Signaling Technologies, 68851), H3K9me2 (EMD Millipore, 05-1249), H3K9me3 (Abcam, AB176916), WIZ (Novus, NBP1-80586), PCNA (Cell Signaling Technologies, 13110), and S33 phosphorylated RPA (Cell Signaling Technologies, 35869). Primary antibodies were selected from different host species to permit recognition by complementary PLA probes. Following primary antibody incubation, slides were washed twice with Duolink Wash Buffer A (included in Duolink® in Situ Wash Buffers, Fluorescence, Sigma-Aldrich, DUO82049) and incubated with species-specific Duolink PLA Probe anti-Mouse PLUS (Sigma-Aldrich, DUO92001) and Duolink PLA Probe anti-Rabbit MINUS (Sigma-Aldrich, DUO92005) probes for 1hr at 37°C in a humidified chamber. Slides were washed twice with Wash Buffer A, and ligation was performed using Duolink Ligation Buffer (included in Duolink® In Situ Detection Reagents Red, Sigma-Aldrich, DUO92008) containing DNA Ligase (included in Duolink® In Situ Detection Reagents Red, Sigma-Aldrich, DUO92008) for 30 min at 37°C. Samples were again washed twice with Wash Buffer A, followed by rolling-circle amplification using Duolink Amplification Buffer (included in Duolink® In Situ Detection Reagents Red, Sigma-Aldrich, DUO92008) containing DNA Polymerase (included in Duolink® In Situ Detection Reagents Red, Sigma-Aldrich, DUO92008) for 100 min at 37°C in the dark inside a humidified chamber. Unless otherwise indicated, PLA probe, ligation, amplification, and wash steps were performed according to the manufacturer’s instructions. Following amplification, cells were incubated in 1μg/mL DAPI (Thermo Fisher Scientific, D1306) along with AF488-conjugated anti-mouse highly cross-adsorbed secondary antibody (Invitrogen, A11029) and Alexa AF647-conjugated anti-rabbit highly cross-adsorbed secondary antibody (Invitrogen, A21244) for 2 hours in the dark at room temperature. Slides were then washed twice with Duolink® Wash Buffer B (included in Duolink® in Situ Wash Buffers, Fluorescence, Sigma-Aldrich, DUO82049), followed by a final wash in diluted Wash Buffer B (0.01X). Coverslips were mounted using ProLong Diamond Antifade Mountant (Invitrogen, P36970) and slides were sealed with nail polish and stored at −20°C and protected from light until imaging. Images were acquired using a Leica SP8 confocal microscope using the 63x and 100x objective. For each experiment, imaging parameters, including exposure time, laser power, gain, and z-stack spacing, were held constant across treatment conditions within antibody groups. Images were acquired as multichannel z-stacks containing DAPI (405nm), EdU (488nm), PLA (594nm), and antibody (647nm) fluorescence channels. Z-stacks were maximum projected prior to quantitative analysis using identical processing parameters across conditions.

Image analysis was performed in FIJI/ImageJ v2.16.0 and R using custom scripts. Briefly, nuclei were segmented using the DAPI channel, and PLA puncta were identified within individual nuclear masks using a consistent intensity and size threshold across samples within an experiment. PLA signal was quantified as the number of PLA puncta per nucleus according to a user-defined threshold which was held constant throughout analysis. Strong EdU fluorescence was used to identify S-phase nuclei prior to PLA quantification, and non-S-phase nuclei were treated as a measurement of background signal. Assay validation controls of PCNA and S33 phosphorylated RPA were included to confirm labeling of nascent DNA and induction of replication stress, respectively. Negative controls included omission of one primary antibody and samples lacking EdU labeling to assess the degree of nonspecific PLA signal. Identical image acquisition and analysis procedures were applied to experimental and control samples.

### Isolation of Proteins on Nascent DNA (iPOND)

Isolation of proteins on nascent DNA (iPOND) was performed according to a previously published protocol with minor modifications^51^. Approximately 50 million cells were used per condition unless otherwise indicated. Cells were maintained at a density of 1-5 × 10^5^ cells/mL and allowed to equilibrate in prewarmed growth medium for at least 30 min prior to labeling. To label nascent DNA, cells were incubated with 10μM 5-ethynyl-2′-deoxyuridine (EdU; Sigma-Aldrich, 900584) for 15 min at 37°C. For replication stress conditions, 1mM hydroxyurea (HU, EMD Millipore, 400046) was added following the 15-minute EdU pulse, and cells were incubated for 1 or 4 hours prior to fixation. For post-replicative DNA controls, cells were pulse-labeled with EdU for 15 min, collected by centrifugation, and washed into prewarmed growth medium containing 10μM thymidine (Sigma-Aldrich, 89270). Cells were then chased with 10μM thymidine for 1 or 4 hours at 37°C prior to fixation. Where indicated, cells were treated with 20μM dWIZ-2 (Selleck Chemicals, E4661) for 4 hours prior to EdU labeling. Cells were fixed by direct addition of 37% formaldehyde (Sigma-Aldrich, 52549) (made fresh) to a final concentration of 1% and incubated for 20 min at room temperature with agitation. Crosslinking was quenched by addition of glycine (Sigma-Aldrich, G8790) to a final concentration of 12.5mM for at least 5 min at room temperature. Cells were collected by centrifugation and washed three times with ice-cold PBS. Cell pellets were either processed immediately or stored at −80°C until further use. For permeabilization, fixed cells were resuspended at 1 × 10^7^ cells/mL in 0.25% Triton-X-100 (Thermo Scientific, 327371000) in PBS and incubated for 30 min at room temperature. Cells were collected by centrifugation, washed once with ice-cold 0.5% BSA (GoldBio, A-420-500) in PBS, followed by one wash with ice-cold PBS, and kept on ice until EdU-labeled nascent DNA was conjugated to biotin using CuAAC click chemistry^60^ as described above with the addition of 2mM hydroxypropyltriazolylmethylamine (THPTA; Vector Laboratories, CCT-1010-100) to inhibit protein oxidation^60^. Briefly, cells were resuspended at 2 × 10^7^ cells/mL in freshly prepared click-reaction buffer containing PBS, 2mM Cu(II)SO4, 2mM THPTA, 10μM biotin picolyl azide, and 10mM sodium ascorbate. Click reactions were allowed to proceed for 1 hour at room temperature with rotation. A no-click control, in which biotin picolyl azide was omitted from the previously described reaction, was processed in parallel to assess nonspecific recovery by the streptavidin pulldown. Following the click reaction, cells were collected by centrifugation, washed once with ice-cold 0.5% BSA in PBS and twice with ice-cold PBS. Cell pellets were resuspended at 1.5 × 10^7^ cells per 100μL in lysis buffer containing 1% sodium dodecyl sulfate (Sigma-Aldrich, L3771) and 50mM Tris-HCl pH 8.0 (Sigma-Aldrich, T5941) supplemented with protease inhibitor (Roche Diagnostics, 82336300). Chromatin was solubilized using a Branson Microtip Sonicator with 3-5 20-second pulses at an output power of 3. Samples were cooled during and between pulses. Samples were centrifuged at maximum speed for 10 minutes at room temperature to remove insoluble material, and the resulting supernatants were passed through a 90-μm cell strainer. Solubilized chromatin was subsequently diluted 1:1 with PBS containing 2× protease inhibitor. Protein concentration was determined using a Pierce BCA assay (Thermo Scientific, 23225) with BSA in water as a standard, and equivalent amounts of protein were used for each pulldown. Approximately 1-3% of each sample was reserved as input. For streptavidin pulldown, streptavidin agarose beads (EMD Millipore, 69203) were prewashed in a 1:1 mixture of lysis buffer and PBS before addition to the solubilized chromatin. For streptavidin agarose pulldowns, 100μL of 50% bead slurry was used per 50 million starting cells. Samples were incubated with streptavidin beads overnight at 4°C with rotation. Following pulldown, beads were collected by centrifugation at 1,800 × g for 3 minutes at room temperature and washed once with 1mL ice-cold lysis buffer for 5 minutes with rotation. Beads were subsequently washed once with 1M NaCl (Sigma-Aldrich, S3014) for 5 minutes, followed by two additional washes with ice-cold lysis buffer. After the final wash, residual buffer was removed using a 31g needle and bound proteins were eluted by addition of one packed-bead volume of 2× LDS sample buffer (GenScript, M00676) containing 5% beta-mercaptoethanol (Sigma-Aldrich, M6250). Samples were heated at 95°C for 30 min to reverse formaldehyde crosslinks and elute bound proteins. Beads were pelleted by centrifugation at maximum speed for 3 min, and the resulting supernatants were collected for analysis by immunoblotting. Input samples were mixed with an equal volume of 2× LDS sample buffer containing β-mercaptoethanol and heated at 95°C for 30 minutes to reverse crosslinks. iPOND eluates and corresponding input samples were analyzed by SDS-PAGE and immunoblotting using antibodies against the indicated proteins and histone modifications as described previously.

### Quantitative Gene Expression Profiling (RT-qPCR)

Pellets were processed using the Zymo RNA Miniprep Kit (Zymo Research, R1054) according to the manufacturer’s protocol. RNA was quantified using a NanoDrop, and 2μg RNA was reverse transcribed into cDNA in a 20μl reaction using the Applied Biosystems cDNA Transcription kit (Thermo Fisher Scientific, 4368814) according to the manufacturer’s instructions. cDNA was then diluted 1:1.5 and 5μl was used for quantitative RT-PCR using the Roche SYBR Green I master mix (Roche, 4707516001) on a Roche LightCycler 480 machine. *PPIB* and *RPLP0* were used as expression controls unless otherwise indicated. The primers used are listed in Table S1.

### Nuclear RNA sequencing

Nuclear RNA-seq was performed with 10 million cells per sample. Briefly, cells were washed twice with PBS and resuspended in 1.6mL of nuclear isolation buffer (1mM Tris-HCl, pH 7.4 (Sigma-Aldrich, T5941), 10mM NaCl (Sigma-Aldrich, S3014), 3mM MgCl2 (Thermo Fisher Scientific, AM9530G), and 0.025% IGEPAL (Sigma-Aldrich, I8896)). Cells were lysed on ice for exactly 5 minutes before rapid spike-in of 6.4mL of wash buffer (1% BSA (GoldBio, A-420-500) in PBS), then pelleted at 600 x g. The supernatant was then aspirated, and the nuclear pellet was washed twice with wash buffer then resuspended in 1mL TRIzol (Aladdin Scientific, T751379). RNA was then extracted via standard TRIzol extraction^61^ and prepped using the Zymo RNA Miniprep Kit (Zymo Research, R1054) according to the manufacturer’s instructions and quantified using a NanoDrop. Library preparation was performed by Plasmidsaurus through poly-A mRNA capture with an oligo-dT primer and reverse transcription for cDNA synthesis. RNA sequencing was performed by Plasmidsaurus through 3’ end counting for transcriptome-wide differential gene expression analysis. Sequencing reads were demultiplexed using BCL Convert v4.3.6 and fqtk v0.3.1. Reads were processed using fastp v0.24.0 to remove poly-X tails and low-quality 3′ sequences, with a minimum Phred quality score of 15 and minimum retained read length of 50 bp. Processed reads were aligned to the human reference genome using STAR v2.7, with non-canonical splice junctions removed. Resulting BAM files were coordinate sorted using samtools v1.21, and PCR and optical duplicates were removed based on unique molecular identifiers (UMIs) using UMICollapse v1.1.0. Alignment quality, strand specificity, and read distribution across genomic features were assessed using RustQC v0.2.1, and sequencing quality metrics were compiled using MultiQC v1.33. Gene-level expression was quantified using featureCounts (Subread v2.1.1) with strand-specific counting and fractional assignment of multi-mapping reads. Reads mapping to annotated exons and 3′ untranslated regions were assigned and grouped by gene ID, with gene biotype and associated metadata obtained from the reference GTF annotation. Differential expression was determined using edgePython v0.2.5. Enriched regulatory motifs were determined using Enrichr^38–40^.

### scRNA-seq data analysis

For data analysis of human CD34^+^-derived erythroid progenitors, FASTQ files were mapped to the human cDNA transcriptome with the 10x v3 whitelist (files available at https://github.com/pachterlab/kallisto-transcriptome-indices/releases) using Kallisto v0.4.6. Samples were demultiplexed and doublets were discarded, then processed using Seurat. Genes expressed in fewer than 3 cells or cells with fewer than 200 distinct transcripts or with a mtRNA percentage of >10% were discarded. For the normal bone marrow reference, raw data from four cell subsets (CD34^+^ CD164^+^, CD34^-^ CD164^hi^, CD34^lo^ CD16^hi^, and CD34^-^ CD164^lo^) were downloaded from GSE117498^62^, and genes expressed in fewer than 3 cells or cells with fewer than 200 distinct transcripts or a mtRNA percentage of >15% were discarded. The reference data was integrated with our scRNA-seq data using the FindTransferAnchors and MapQuery functions in Seurat, and projections of this scRNA-seq data to the reference map were used for subsequent analyses. Pseudotimes for cells along the erythroid trajectory were calculated using the Slingshot package in R (v2.10.0) and expression of specific genes was plotted using the geom_smooth function.

### ATAC-seq

ATAC-seq was performed using the Active Motif ATAC-seq Kit (Active Motif, #52150) according to the manufacturer’s protocol. HUDEP-2 HBG-mCherry cells were sorted based on their mCherry expression into ice-cold media using the FACSAria as described previously, then washed in PBS twice and counted. Nuclei were isolated from 100,000 cells using 100μL hypotonic buffer (10mM Tris pH 7.4, 10mM NaCl, 3mM MgCl2, 0.1% NP40) to generate two independent transposition reactions. Nuclei were split in half and treated with 2.5μl Tn5 transposase for 30 minutes at 37°C. DNA from transposed nuclei was then isolated and PCR amplified using barcoded Nextera primers (Illumina). Library QC was carried out using High Sensitivity DNA Bioanalyzer assay (Agilent, 5067-4626) and Qubit measurement and sequenced using paired-end sequencing (PE50) on the Illumina HiSeq 2500 platform.

Human data was trimmed using Cutadapt (v3.4) (with parameters m 20 -q 20 -a CTGTCTCTTATA -A CTGTCTCTTATA), then aligned to the hg38 genome with Bowtie2 v2.4.2 (with parameters -- dovetail -I 10 -X 1000 --very-sensitive). Reads in the mitochondrial genome and unannotated chromosomes as well as the UCSC blacklist were removed, then duplicates were removed with samtools markdup (v1.1.2), and the insert size distribution was calculated using Picard CollectInsertSizeMetrics (v2.25.2) and validated to contain nucleosomal peaks. The BED file was then constructed with each mate pair as a separate read using bedtools bamtobed (v2.3.0), and peaks were called using MACS2 callpeak (v2.2.7.1) (with parameters --nomodel --shift −100 -- extsize 200 --keep-dup all --call-summits). Note that the MACS2 command shifts each read by 100 base pairs towards the 5’ end and extends the read length to 200 base pairs, thus treating the 5’ end of the original read (the Tn5 cut site) as the center of a new “fragment” that is then used for peak calling and other downstream analysis. To normalize tracks for visualization, reads overlapping shared peaks were counted for each BAM file using featureCounts (with parameters -M -O) and normalization factors were calculated from these counts using the calcNormFactors function in the edgeR package (v3.32.1) in R. Visualization was performed using the bedGraphToBigWig utility. To identify changes in motif accessibility, ChromVAR^63^ was used. For downstream analysis, peaks from all samples from the same experiment were merged as follows. Peak summits (called using MACS2) were extended by 25bp in each direction, then only peaks that were present in both replicates for each condition (overlapping by at least 50%) were retained. Peak lists were then merged across samples, merging peaks that were within 50bp of each other. ATAC-seq reads were then counted against this master peak list using featureCounts (-p -O -M --fraction). Differential ATAC-seq peaks were calculated using edgeR with a log_2_ fold change threshold of 1 and an adjusted p-value of 0.05. Peaks were annotated to genomic features using ChIPseeker^64^.

### Transcription factor motif, pathway, and gene-set enrichment analysis

Enriched regulatory motifs and pathways were determined using Enrichr^38–40^. Gene set enrichment analysis (GSEA) was conducted using GSEA_v4.4.0. To generate the GATA1 target gene signature, we compiled the 150 most significantly downregulated genes following GATA1 knockdown (reanalyzed from GSE214811^41^, including all timepoints). GSEA was performed using rank-ordered log_2_(Fold Change) values and the GSEA “Preranked” option.

### Reanalysis of Published ChIP-seq data

BigWig files were downloaded from GSE298130^65^ and visualized using IGV 2.19.5.

### Statistical Analysis

All experiments were performed with at least three independent biological replicates unless otherwise indicated. Data are presented as mean ± SD unless otherwise indicated. Statistical analyses were conducted using appropriate parametric or nonparametric tests as specified in the figure legends.

## Notes

https://github.com/oaksgil/Oaks-Do-RS-redistributes-G9a

