## Supplemental Figures and Legends for "DNA replication stress redistributes the histone methyltransferase G9a and activates lineage-specific gene expression"

### Supplemental Material

#### **Figure S1: Inhibitors of DNA replication increase gamma-globin expression in erythroid cells. Related to Figure 1.**

**(A)** Dose-response curve of percent hemoglobin F (HbF)-expressing (red, left axis) and cell viability (black, right axis) in HUDEP-2 cells treated with increasing concentrations of camptothecin. Data represent mean ± SEM (n=3).

**(B)** Representative immunoblot of HUDEP-2 cells treated with increasing doses of camptothecin for p-Chk1 and p-Chk2 expression, with Vinculin included as a loading control.

**(C)** Representative flow cytometry gating strategy for primary human CD34<sup>+</sup> hematopoietic stem and progenitor cells (HSPC)-derived erythroid progenitors using CD235a and CD71 staining to define erythroid populations, with percentages of each gated population indicated.

**(D)** Dose-response curve of fetal hemoglobin induction (HbF, red) in CD34<sup>+</sup> HSPCs treated with increasing concentrations of decitabine. Data represent mean ± SEM (n=3).

**(E)** Dose-response curve of p-Chk1 staining as measured by intracellular flow cytometry staining in CD34<sup>+</sup> HSPCs treated with increasing concentrations of hydroxyurea (HU), aphidicolin, and decitabine. Data represent mean ± SEM (n=3).

**(F)** Cell surface marker staining of erythroid markers CD235a and CD71 in HUDEP-2 cells treated with the indicated doses of HU. Bars represent the mean percentage of cells in each population (n=3).

**(G)** Dot plots showing expression of erythroid-associated genes (*HBB*, *HBA1*, *GATA1*, *BCL11A*, and *TMCC2*) in erythroid clusters 1–3 (see Figure 1G) of CD34<sup>+</sup> HSPCs treated with vehicle (Veh), HU, or aphidicolin (APH) from days 7–14 of erythroid differentiation. Dot size represents the percentage of cells expressing each transcript, and color indicates scaled average transcript expression.

**Figure S2: A live cell reporter for HbF allows investigation of the dynamics of fetal hemoglobin activation and repression. Related to Figure 1.**

**(A)** Schematic of the HBG2 reporter in HUDEP-2 cells, showing insertion of a P2A-mCherry cassette downstream of *HBG2* to track *HBG2* expression (HBG-mCherry cells). Also shown is PCR validation of correct mCherry insertion into the HBG2 locus with expected fragment sizes corresponding to that indicated on the schematic.

**(B)** Representative flow cytometry plots (top) showing induction of mCherry following treatment with hydroxyurea (HU) or decitabine (DEC). Quantification (bottom) shows the percentage of mCherry+ cells at different doses of these compounds after 24, 48, or 72 hr treatment. Data represent mean ± SEM (n=3).

**(C)** Correlation of mCherry reporter signal with HbF protein expression. Cells were sorted into mCherry-low (A cells) and mCherry-high (F cells) populations at baseline and analyzed for intracellular HbF staining after HU treatment. Representative flow cytometry plots and percentages of HbF+ cells are shown.

**(D)** Gene expression analysis of globin genes across multiple HUDEP2 HBG-mCherry cell populations including the bulk parental cells (Total), and mCherry-low, mCherry-medium, and mCherry-high cells as indicated. Left: *HBG* expression normalized to *AHSP* and *RPS18*. Middle: *HBB* expression normalized to *AHSP* and *RPS18*. Right: fraction of HBG relative to total globin gene expression ($HBG/(HBG+HBB)$). Data represent mean ± SEM (n=3).

**(E)** Schematic of HUDEP-2 HBG-mCherry single cell clones, with baseline mCherry+ cell percentage denoted as a proxy for baseline HbF.

**(F)** Dose-response curve of mCherry+ percentage (red, left axis) and cell viability as measured by flow cytometry (black, right axis) for HUDEP-2 single cell clones described in (E) treated with increasing concentrations of HU (top) and APH (bottom) as indicated. Data represent mean ± SEM (n=3).

**(G)** Schematic showing experimental design to assess stability and reversibility of HbF activation. Reporter cells were pretreated for 48 hr with Veh, HU, or DEC, and then were sorted into A (low mCherry) and F (high mCherry) cell populations. All populations were then allowed to recover in Veh or HU and the percent of A and F cells was reanalyzed after three days.

**(H)** Plots showing the percent of F (mCherry+) cells after sorting A cell populations (left) or F cell populations (right) followed by three days of exposure to either Veh or HU. The percent of F cells immediately post-sorting is shown for both sorted populations.

**(I)** Schematic showing experimental workflow to assess heritable, non-genetic memory in HUDEP-2 cells. Single A (mCherry low) cells were sorted, expanded into colonies, split into three sister colonies, and treated with either vehicle (one replicate) or HU (two replicates).

**(J)** Correlation analysis of percent F cells per colony between two independent HU treatment replicates derived from the same single cell population. Each point represents a single colony, and point color indicates baseline (untreated) percent F cells. The regression line and 95% confidence interval are shown, along with Pearson correlation coefficient and p-value.

**(K)** Boxplot of the average percent F cells for the two HU treated sister colonies, binned by the percentage of F cells in the vehicle treated colony. P-value was generated using a Mann-Whitney U test.

**Figure S3: Early HbF induction by replication stress is independent of replication stress signaling or HRI. Related to Figure 1.**

- **(A)** Representative immunoblot of parental and HRI knockdown HBG-mCherry HUDEP-2 cells treated with vehicle (Veh), 50 μM hydroxyurea (HU), 6.25 μM gemcitabine (GEM), or 10 μM pomalidomide (POM) for 72 hr. The band corresponding to HRI is indicated with an arrow.
- **(B)** Dose-response curve of fetal hemoglobin induction (HbF, red, left axis) and cell viability (black, right axis) in parental (WT, left) and HRI knockdown (shHRI, right) HUDEP-2 cells treated with increasing concentrations of decitabine, camptothecin, and cytarabine as indicated (n=3).
- **(C)** Representative immunoblot of HUDEP-2 cells pretreated for 24 hr with HU and then treated with increasing doses of AZ20 for 4 hr assessing expression of total CHEK1 (Chk1) and total CHEK2 (Chk2) as well as phospho-CHEK1 (p-Chk1) and phospho-CHEK2 (p-Chk2), with PCNA presented as a loading control.
- **(D)** Percent viability as measured by flow cytometry for HUDEP-2 HBG-mCherry cells treated with Veh or two different doses of AZ20 in addition to Veh or the indicated dose of HU or aphidicolin (APH) for 12 or 24 hr as indicated.
- **(E)** Percent mCherry+ HUDEP-2 HBG-mCherry cells following treatment with vehicle or two different doses of AZ20 in addition to Veh or the indicated dose of HU or APH for 12 or 24 hr as indicated. The dashed line indicates the baseline percentage of mCherry+ cells for the HUDEP-2 cells used in this experiment.

**Figure S4: H3K9me2, G9a, and WIZ are reduced at the gamma-globin locus and genome-wide. Related to Figure 2.**

- **(A)** Genome browser tracks showing CUT&RUN signal for the indicated histone marks at the β-globin locus in HUDEP-2 HBG-mCherry Clone C (see Figure S2E, ~20-30% mCherry+ cells at baseline), normalized to counts per million (CPM). Tracks display H3K9me2, H3K9me3, H3K9ac,

H3K27ac, and IgG control after 24 hr treatment with vehicle (Veh, blue) or hydroxyurea (HU, red). Highlighted regions indicate the γ-globin genes (*HBG1/2*) and locus control region (LCR).

**(B)** Genome browser tracks showing CUT&RUN signal for the indicated histone marks at the β-globin locus in HUDEP-2 Clone A (see Figure S2E, ~5% mCherry+ cells at baseline), normalized to mouse spike-in controls. Tracks show H3K9me2, H3K9me3, H3K9ac, H3K27ac, H3K27me3, and IgG control after 24 hr treatment with vehicle (Veh, blue) or hydroxyurea (HU, red). Highlighted regions indicate the γ-globin genes (*HBG1/2*) and locus control region (LCR).

**(C)** CUT&RUN profiles for Clone C showing the same histone marks and conditions as in (A) but normalized to mouse spike-in controls instead of counts per million. Highlighted regions indicate the γ-globin genes (*HBG1/2*) and locus control region (LCR).

**(D)** Corepressor CUT&RUN profiles for Clone A showing G9a, WIZ, and IgG under Veh (blue) and HU (red) conditions normalized to mouse spike-in controls instead of counts per million. Highlighted regions indicate the γ-globin genes (*HBG1/2*) and locus control region (LCR).

**(E)** Genome-wide differential analysis of H3K9me2 signal upon HU treatment in Clone C after normalizing to mouse spike-in (top) or counts per million (bottom). Volcano plots show $\log_2$ of the fold change of HU vs. Veh versus their statistical significance. The number of statistically significant regions for each normalization method is indicated.

**(F)** Dose-dependent effects of UNC-0646 on mCherry induction (% mCherry+ cells, left axis, red) and cell viability (% DAPI- cells, right axis, black) in HUDEP-2 clones B (~10% mCherry+ cells at baseline) and C (~20-30% mCherry+ cells) after 24 hr (n=3).

**(G)** Genome-wide changes in corepressor and histone mark peaks following HU treatment in Clone A. Volcano plots depict $\log_2$ fold change in CUT&RUN signal versus $-\log_{10}$ adjusted p-value for G9a, WIZ, H3K9me2, H3K9me3, and H3K9ac at their respective peak sets, with the number of significantly up- or downregulated regions indicated.

- **(H)** Genome-wide changes in CUT&RUN signal for various histone marks at WIZ-only (red), G9a-only (blue), and WIZ/G9a-cobound peaks (black). Profiles are shown for multiple histone marks (H3K4me1, H3K4me3, H3K27me3, H3K27ac, H3K9me3, H3K9ac) under Veh (columns 1 and 3) and HU (columns 2 and 4) conditions as indicated.
- **(I)** Violin plots showing $\log_2$ fold changes in H3K9me2 (top) or H3K9me3 (bottom) signal upon HU treatment compared to Veh as a function of distance from the nearest WIZ peak.
- **(J)** Overlap between BCL11A peaks and WIZ/G9a co-bound peaks. The number of peaks in each region is indicated.
- **(K)** Aggregate BCL11A CUT&RUN signal at WIZ-only (red), G9a-only (blue), and WIZ/G9a-cobound peaks (black) after Veh (left) and HU (right) treatment.

**Figure S5: Further analysis of changes in chromatin accessibility, H3K9me2, and H3K27me3 with HU treatment. Related to Figure 2.**

- **(A)** Representative genome browser tracks showing ATAC-seq signal in vehicle-treated (Veh, top) and hydroxyurea-treated (HU, bottom) conditions for A cells (low % mCherry+ cells) and F cells (high % mCherry+ cells) across the globin locus.
- **(B)** Bar plot summarizing the number of differentially accessible regions (ATAC-seq peaks with $\log_2FC \geq 1$ and adjusted $p < 0.05$) for the indicated comparisons (HU F vs HU A; Veh F vs Veh A; HU A vs Veh A; HU F vs Veh F), showing gains (Up) and losses (Down) of accessibility.
- **(C)** Volcano plots showing changes in ChromVAR transcription factor motif accessibility scores for HU versus vehicle-treated A cells (top) and vehicle-treated F cells versus vehicle-treated A cells (bottom). Selected significantly enriched motifs are labeled.
- **(D)** Average profiles and heatmaps for H3K9me2, H3K9me3, H3K9ac, H3K27me3, and H3K27ac CUT&RUN signal at ATAC-seq peaks (open chromatin regions, OCRs) within H3K9me2+ regions

following both vehicle and HU treatment. Data are stratified by peaks that lose H3K9me2 with HU treatment, have unchanged H3K9me2 after HU treatment, or have increased H3K9me2 following HU treatment as indicated.

**(E)** Representative genome tracks showing CUT&RUN signal for the indicated histone marks, corepressors or ATAC-seq signal at representative loci (*KIT*, *TRAF6*, *RAG1*, *RAG2*) associated with loss of H3K9me2 in response to HU treatment.

**(F)** Volcano plots showing differential H3K27me3 CUT&RUN signal between Veh and HU at H3K27me3 peaks (top) and at WIZ-bound peaks (bottom). Numbers of significantly increased or decreased regions are indicated.

**(G)** Average signal profiles of histone modifications (H3K4me1, H3K4me3, H3K27me3, H3K27ac, H3K9me2, H3K9me3, H3K9ac) and ATAC-seq centered on OCRs within H3K27me3-marked regions. Red represents regions that lose H3K27me3 in response to HU treatment, black represents regions that have unchanged H3K27me3 in response to HU, and blue represents regions that have increased H3K27me3 in response to HU.

**(H)** Average profiles of G9a, WIZ, and BCL11A occupancy (baseline CUT&RUN signal) at OCRs located within H3K9me2+ or H3K27me3+ regions, stratified by whether H3K9me2 or H3K27me3 decreases, remains unchanged, or increases upon HU treatment.

**(I)** Baseline CUT&RUN signal profiles for H3K9me2 and H3K27me3 across regions that lose H3K9me2 (left) or H3K27me3 (right) following HU treatment.

**(J)** Enriched categories within genes adjacent to OCRs that lose H3K27me3 following treatment with HU.

**(K)** Genome browser tracks showing CUT&RUN tracks for H3K9me2, H3K27me3, G9a, WIZ, H3K4me1, H3K4me3, and ATAC-seq signal under Veh (blue) and HU (red) conditions at representative loci (*DLL1*, *FOXA1*, *NKX3-2*, *SOX4*) that lose H3K27me3.

**(L)** Transcription factor motifs enriched within OCRs that lose H3K27me3 upon HU treatment, with corresponding HOMER p-values and associated transcription factors indicated.

**Figure S6: Redistribution of G9a, WIZ, H3K9me2 contributes to GATA1/2 regulated gene activation. Related to Figure 3.**

**(A)** Venn diagram showing genes associated with WIZ/G9a co-bound peaks (blue) and genes associated with ATAC-seq peaks (OCRs, red) in HUDEP-2 cells. The percent overlap is indicated.

**(B)** Transcription factor motif enrichment analysis at BCL11A-bound peaks. P-values indicate significance of enrichment compared to a background set of peaks as calculated by HOMER.

**(C)** Batch-corrected PCA analysis of nuclear RNA-seq data from untreated cells and cells treated for 24 hr with hydroxyurea (HU), UNC-0646, or dWIZ-2. HU nuclear RNA-seq samples are indicated by circles, while UNC-0646 and dWIZ-2 RNA-seq samples are indicated by triangles.

**(D)** Transcription factor regulatory motif analysis of genes showing significantly increased expression ($\log_2$ fold change > 0.1; adjusted p-value < 0.05) after both UNC-0646 and dWIZ-2 treatment for 24 hr.

**(E)** Scatterplot of genes upregulated by HU treatment, with $\log_2$ fold change of HU relative to vehicle treatment (Veh) shown on the X axis and $\log_2$ fold change of dWIZ-2 relative to Veh on the Y axis. An upregulated subset of the GATA1 target genes used for GSEA in Figure 3G are labeled and Spearman's correlation coefficient and associated p-value are indicated.

**(F)** Venn diagram showing genes upregulated ($\log_2$ fold change > 0.1; adjusted p-value < 0.05) by HU treatment and genes upregulated ($\log_2$ fold change > 0.1; adjusted p-value < 0.05) upon both UNC-0646 and dWIZ-2 treatment. P-value indicates the significance of overlap by Fisher's exact test.

**Figure S7: Corepressor expression does not change at the RNA level following HU treatment and evidence for G9a overexpression in K562 cells. Related to Figure 4.**

**(A)** Bar plots showing log<sub>2</sub> fold change in gene expression of *EHMT2*, *EHMT1*, *WIZ*, *ZNF644*, and *BCL11A* relative to vehicle after 1 hr, 4 hr, and 24 hr HU treatment. Significance ($p < 0.00001$) is indicated by an asterisk ($n=3$ for 1 hr and 4 hr conditions, and $n=4$ for vehicle and 24hr conditions).

**(B)** Representative immunoblot for FLAG and G9a expression in parental K562 cells, K562 cells transduced with a FLAG-G9a overexpression construct prior to puromycin selection, and transduced K562 cells selected with puromycin for 72 hr and maintained in normal culture conditions for at least 48 hr. β-actin is shown as a loading control.

**(C)** Quantification of the immunoblot shown in (B). FLAG and G9a signal is normalized to the corresponding β-actin loading control.

**Figure S8: PLA controls, including evidence dWIZ-2 reduces colocalization of G9a, WIZ, and H3K9me2 with replication forks. Related to Figure 5.**

**(A)** Representative images of DAPI staining (left) and biotin staining following CuAAC click chemistry conjugation<sup>60</sup> of biotin to incorporated EdU (right). Representative cells designated as EdU-positive, and therefore in S phase, are indicated with dashed circles.

**(B)** Representative immunofluorescence images of PLA foci for PCNA and S33 phosphorylated RPA (p-RPA) in proximity to nascent DNA (labeled by EdU) using a proximity ligation assay (PLA). PLA signals (red) are shown for vehicle-treated (Veh), hydroxyurea-treated (HU), and post-replication (post-rep DNA) conditions as described in Figure 5A. Nuclei are counterstained with DAPI (blue). S-phase cells, determined by the presence of EdU-labeled DNA as shown in (A), are indicated with white dashed circles.

**(C)** Quantification of PLA foci assessing proximity of EdU with PCNA or p-RPA from the PLA shown in (B). Data are presented as box-and-whisker plots showing median and interquartile range. Statistical significance was calculated using a Wilcoxon test with Benjamini-Hochberg p-value correction (\*\*p < 0.01, \*\*\*p < 0.001, \*\*\*\*p < 0.0001; ns, not significant). The number of nuclei assessed for each condition is indicated.

**(D)** Quantification of PLA foci assessing proximity of EdU with G9a, H3K9me2, WIZ, H3K9me3, p-RPA, or PCNA under Veh, HU, and Post-rep DNA conditions in Clone C (~20-30% mCherry+ cells at baseline). The number of nuclei assessed for each condition is indicated.

**(E)** Quantification of WIZ-EdU PLA foci under vehicle and dWIZ-2-treated conditions. The number of nuclei assessed for each condition is indicated.

Figure S1: Related to Figure 1.

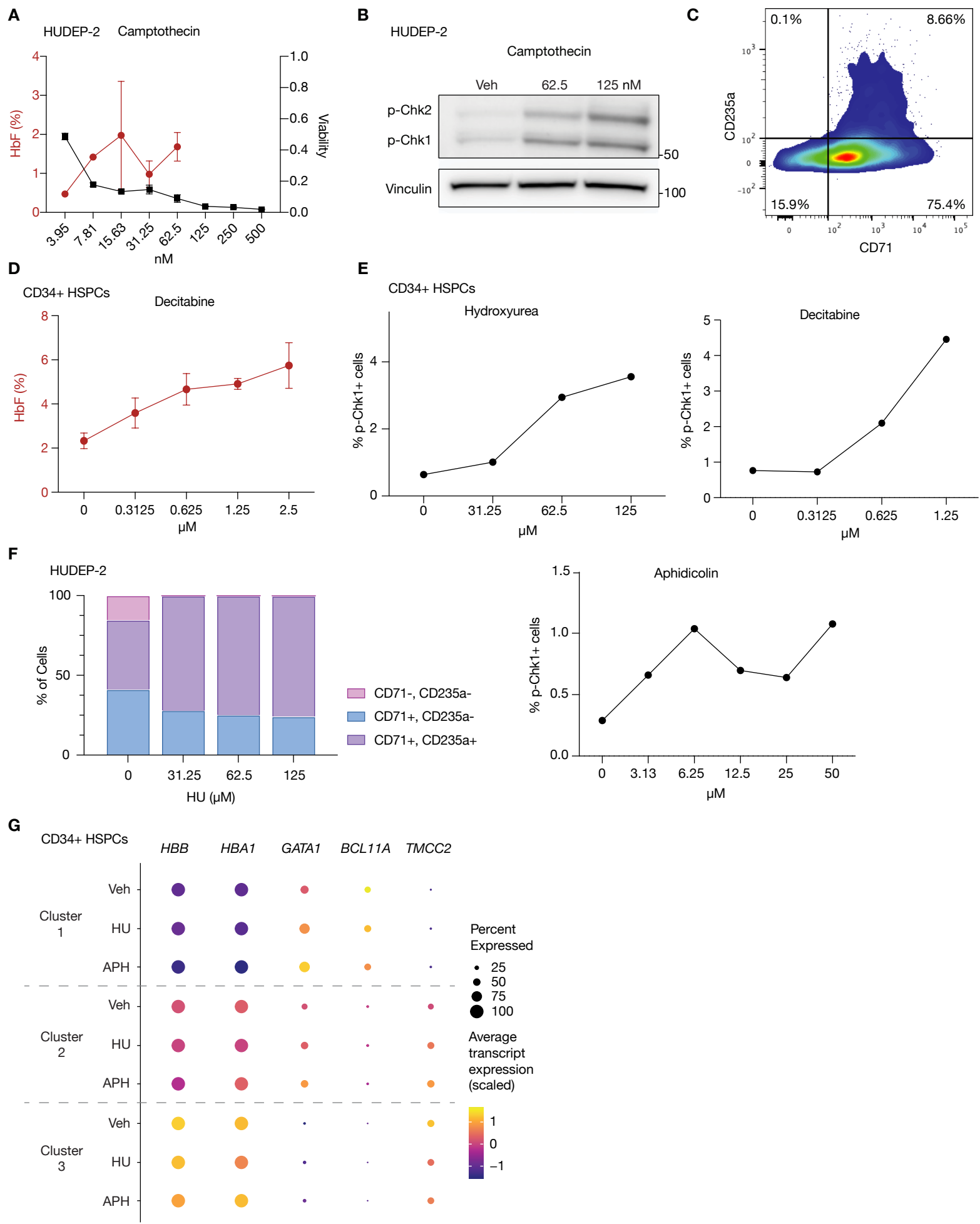

**Figure S2**

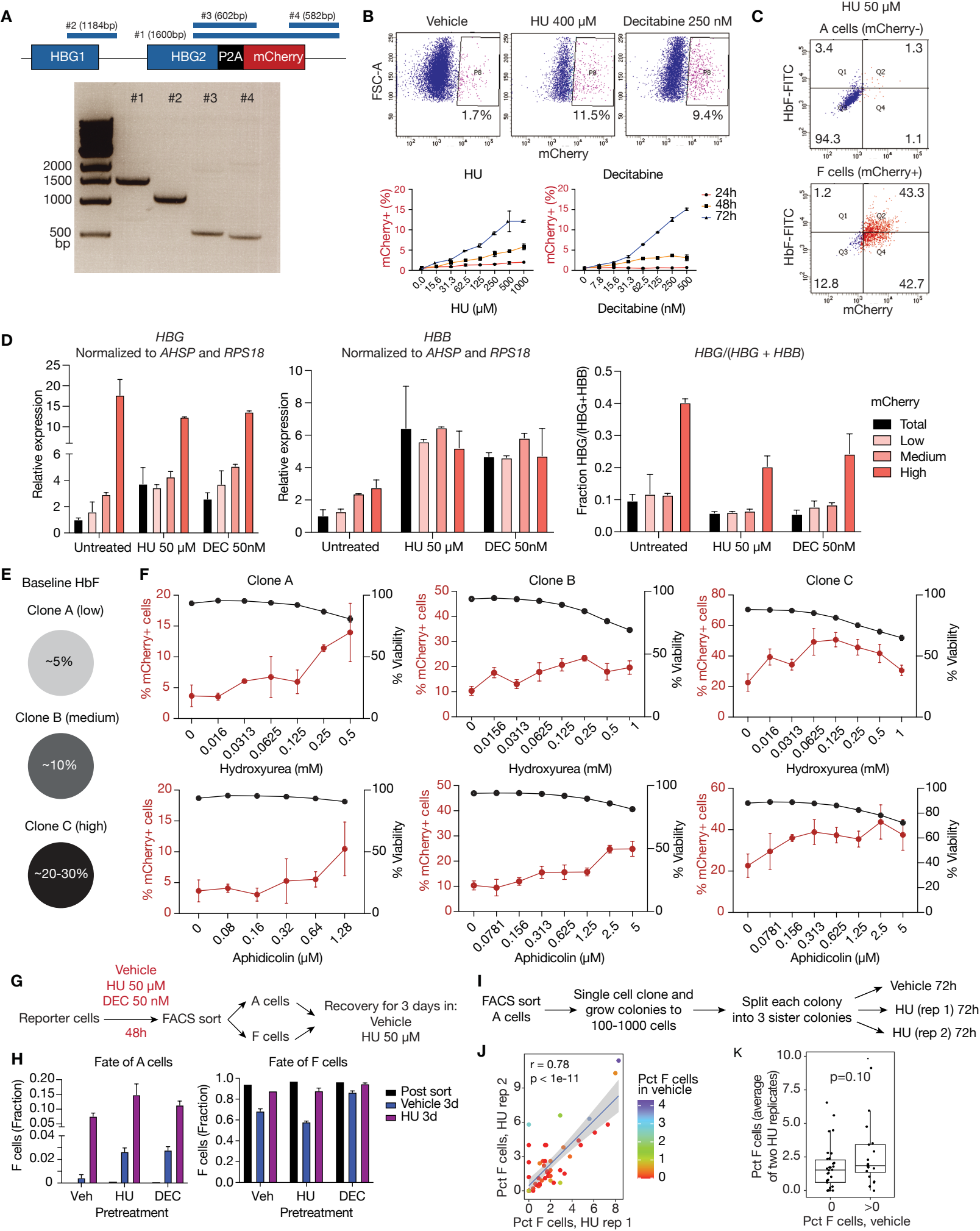

**Figure S3**

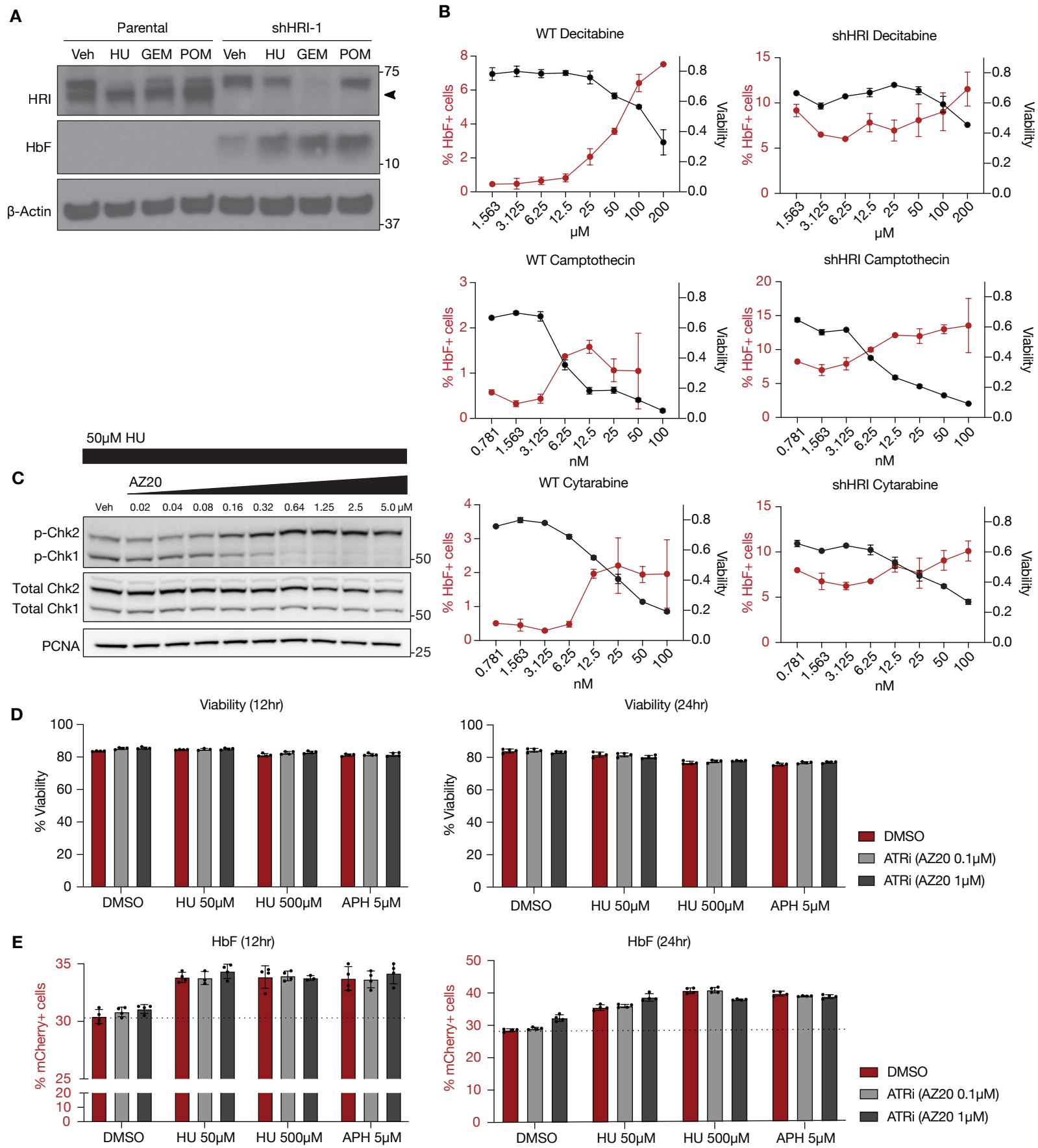

**Figure S4: Related to Figure 2.**

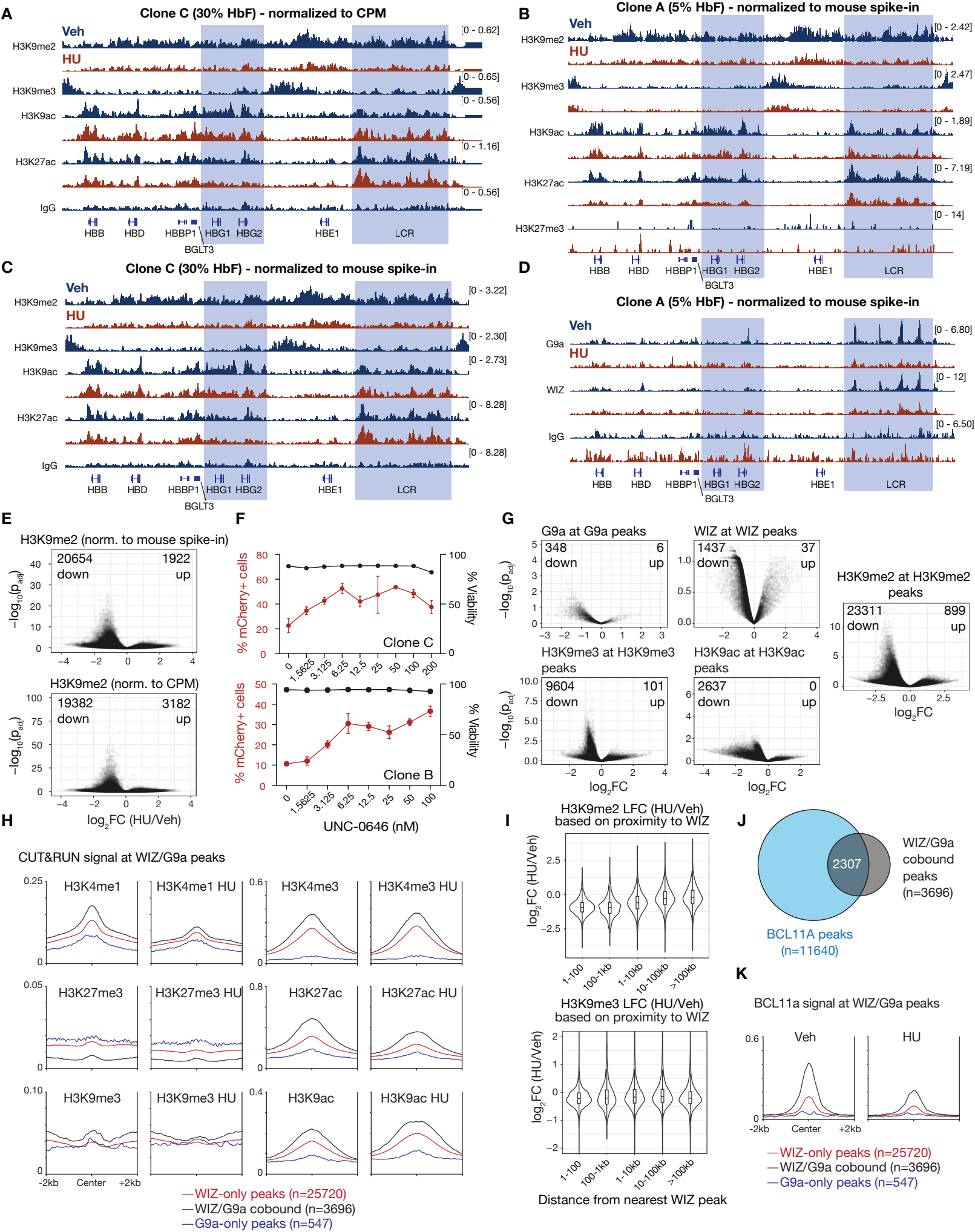

**Figure S5: Related to Figure 2.**

**A**

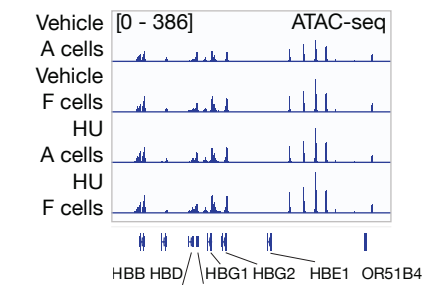

**B**

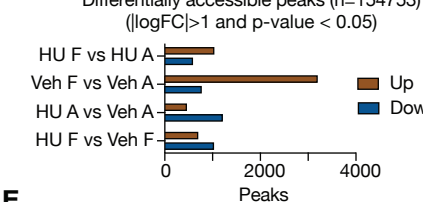

**E**

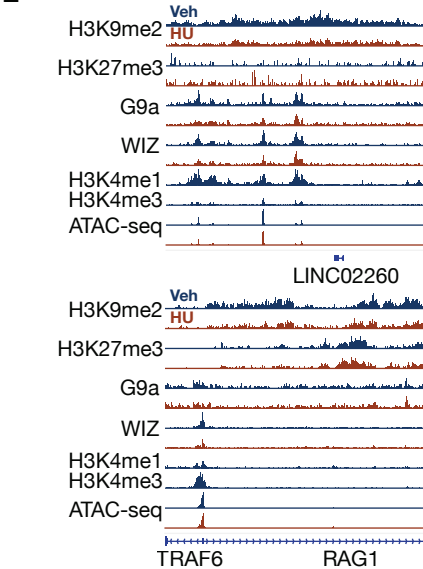

**F**

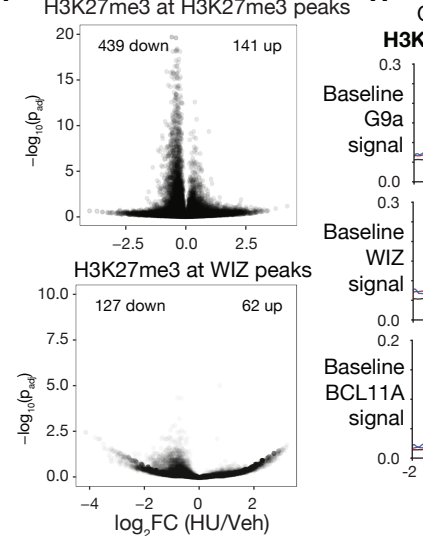

**K**

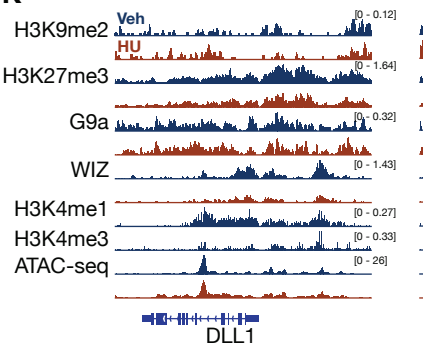

**C**

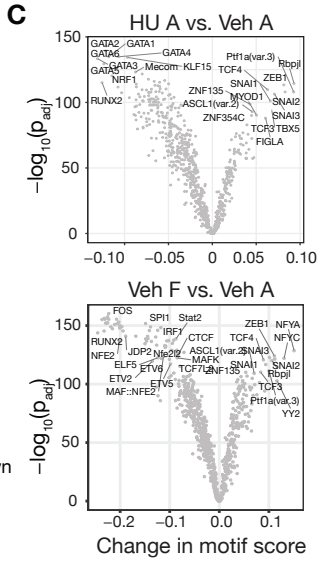

**H**

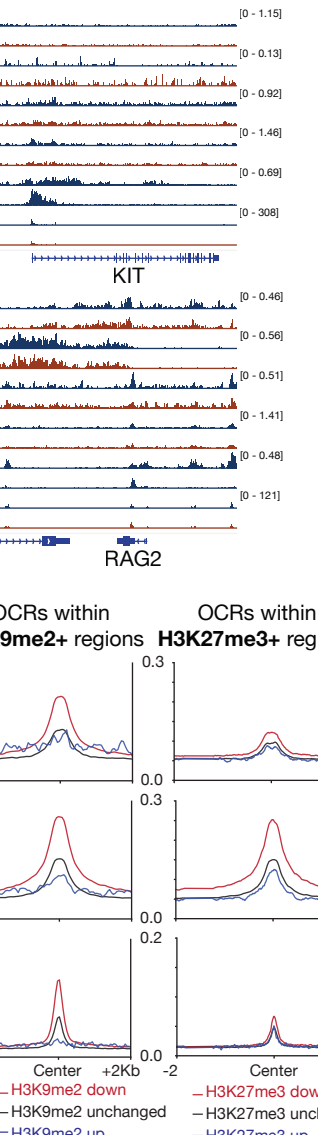

**L**

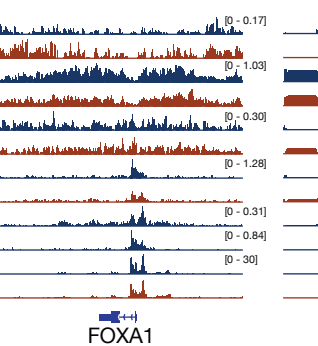

**D**

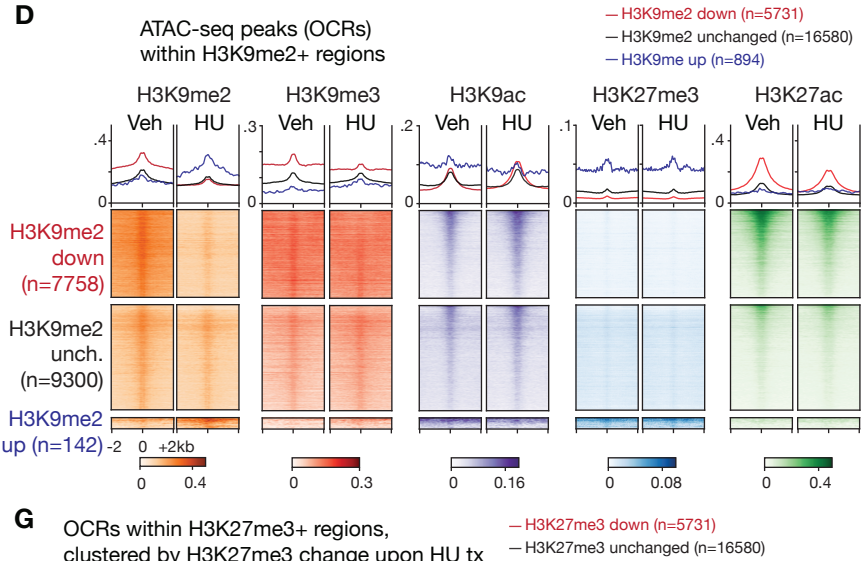

**G**

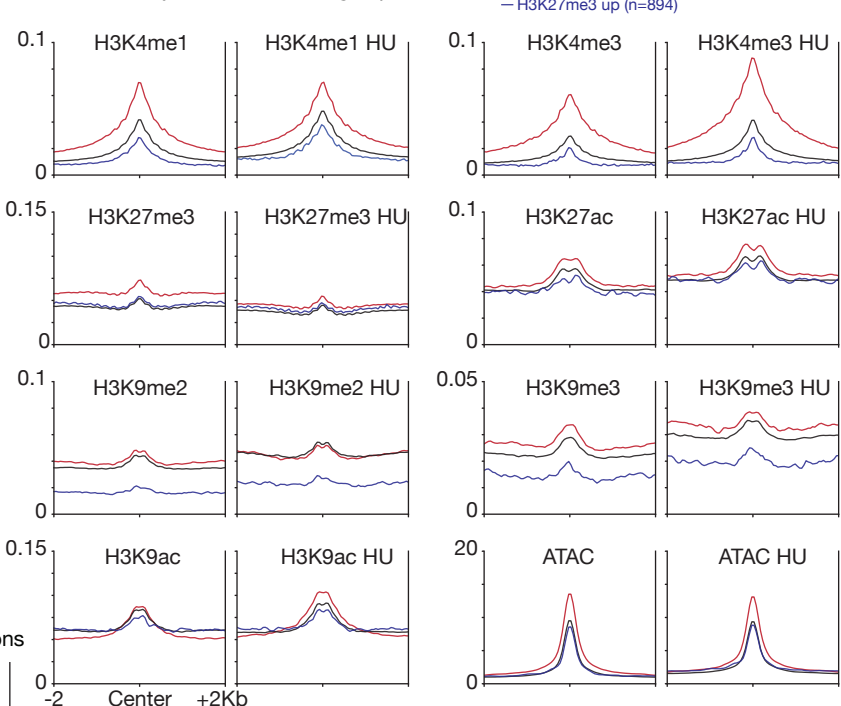

**I**

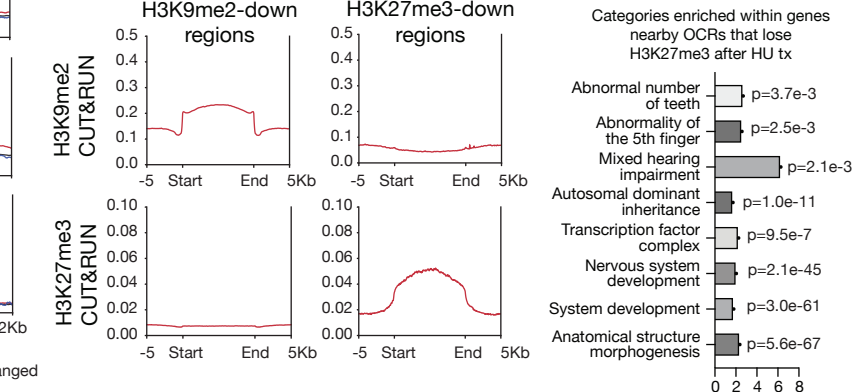

**J**

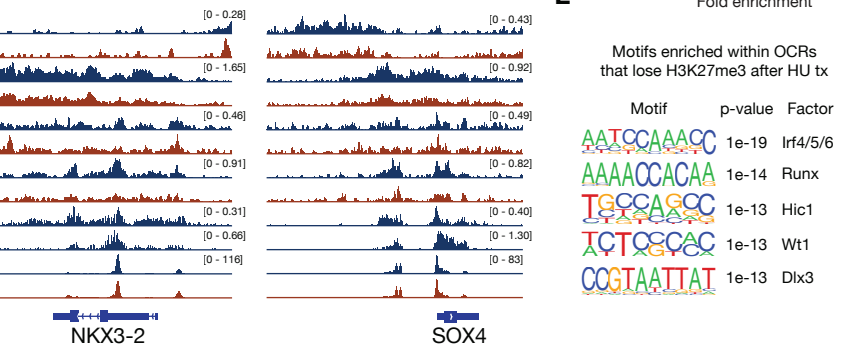

Figure S6: Related to Figure 3.

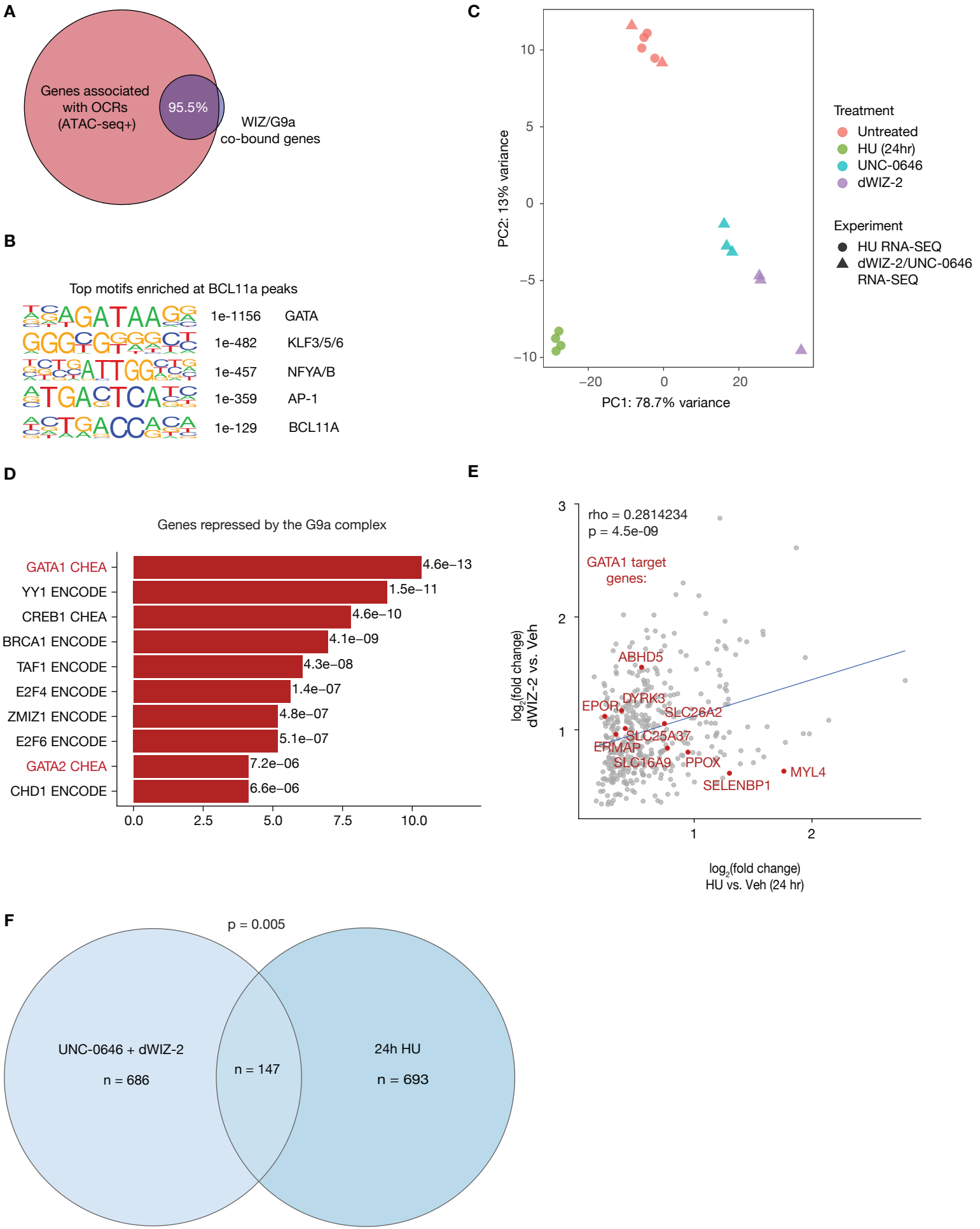

**Figure S7: Related to Figure 4.**

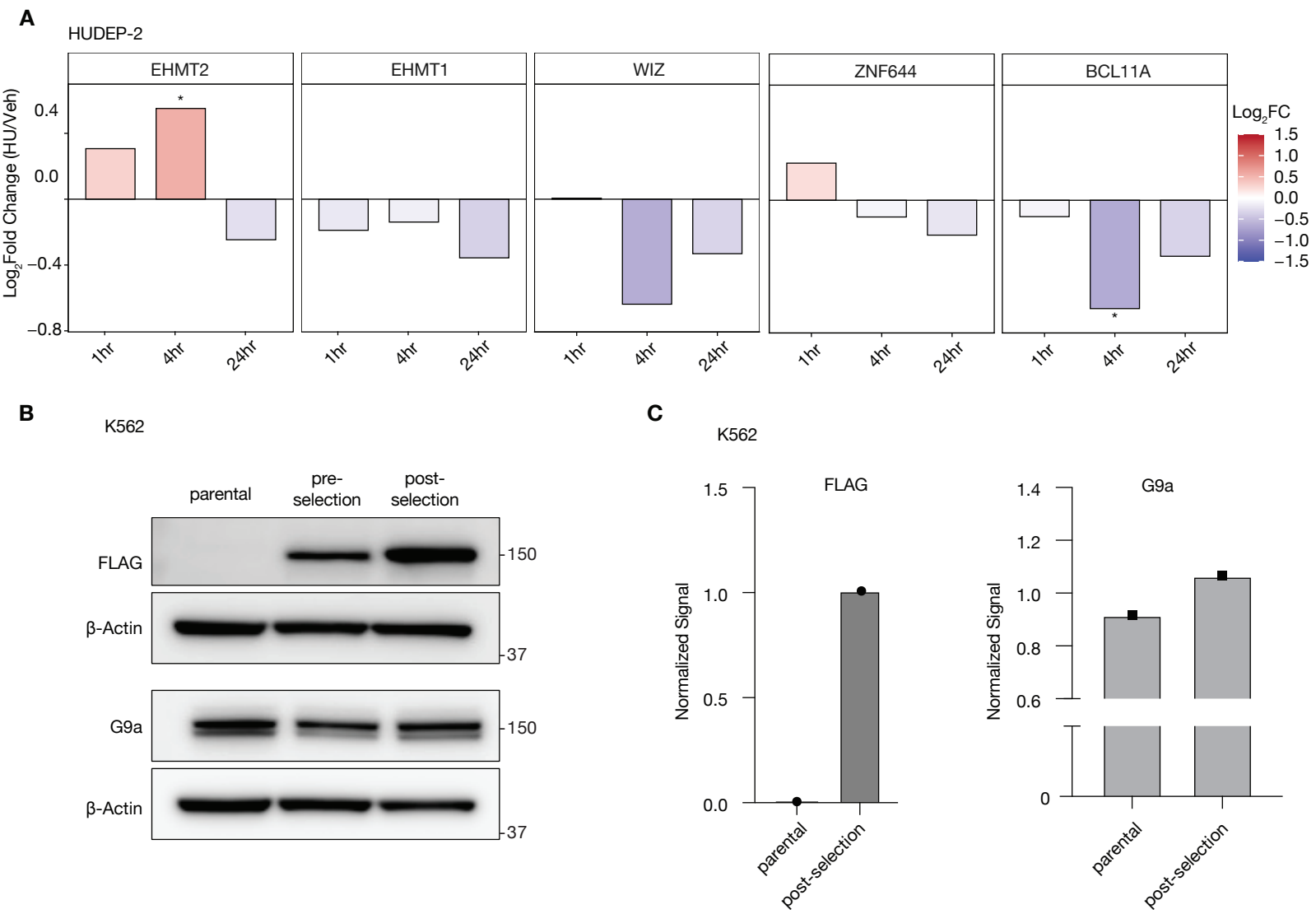

**Figure S8: Related to Figure 5.**

**A** Representative biotin staining with representative selection of S-phase cells

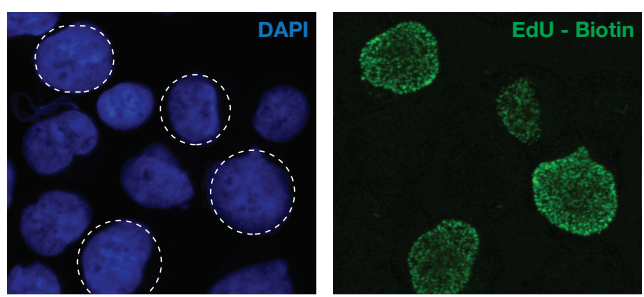

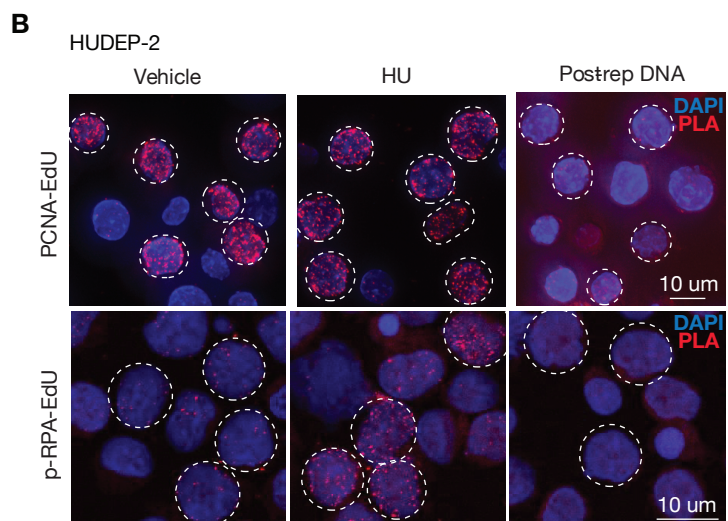

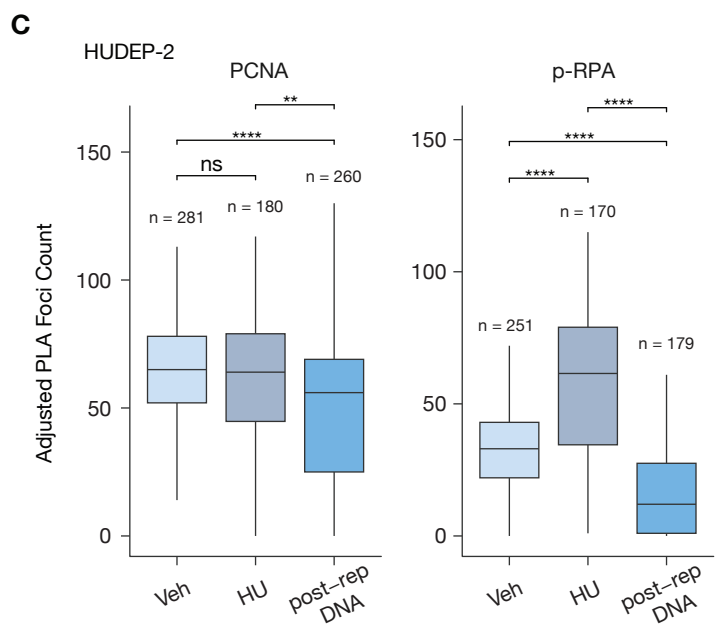

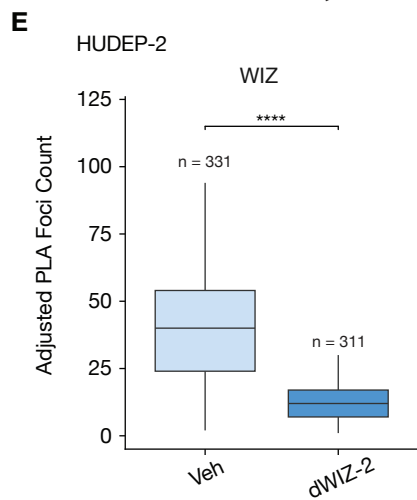

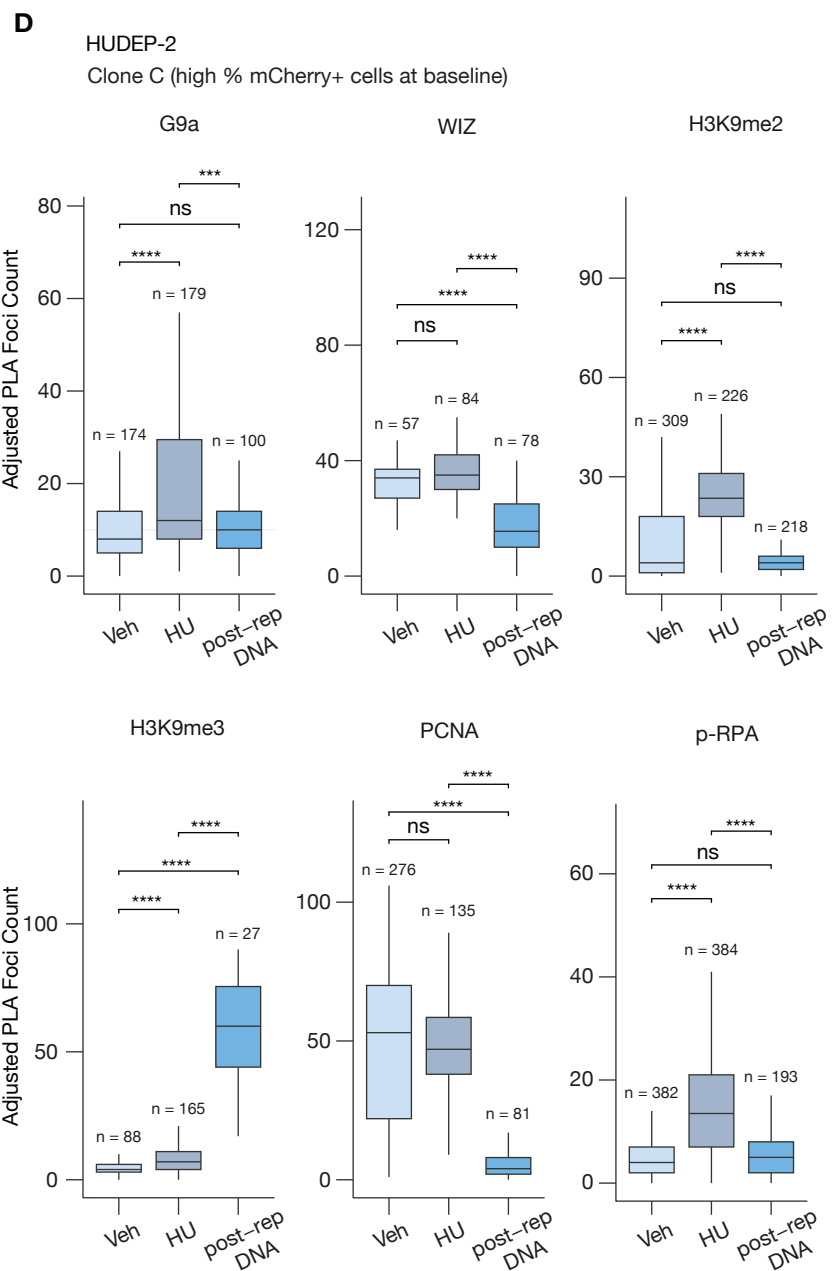
